# Genomes of a major nosocomial pathogen *Enterococcus faecium* are shaped by adaptive evolution of the chromosome and plasmidome

**DOI:** 10.1101/530725

**Authors:** S Arredondo-Alonso, J Top, AC Schürch, A McNally, S Puranen, M Pesonen, J Pensar, P Marttinen, JC Braat, MRC Rogers, W van Schaik, S Kaski, J Corander, RJL Willems

**Affiliations:** Department of Medical Microbiology, University Medical Center Utrecht, Utrecht, The Netherlands; Institute of Microbiology and Infection, University of Birmingham, Birmingham, United Kingdom; Department of Computer Science, Aalto University, FI-00076 Espoo, Finland; Department of Mathematics and Statistics, Helsinki Institute of Information Technology (HIIT), FI- 00014 University of Helsinki, Finland; Pathogen Genomics, Wellcome Trust Sanger Institute, Cambridge CB10 1SA, UK; Department of Biostatistics, University of Oslo, 0317 Oslo, Norway

**Keywords:** *Enterococcus faecium*, adaptive evolution, short-and long-read sequencing, machine-learning, plasmidome, genome-wide co-evolution analysis

## Abstract

*Enterococcus faecium* is a gut commensal of many mammals but is also recognized as a major nosocomial human pathogen, as it is listed on the WHO global priority list of multi-drug resistant organisms. Previous research has suggested that nosocomial strains have multiple zoonotic origins and are only distantly related to those involved in human commensal colonization. Here we present the first comprehensive population-wide joint genomic analysis of hospital, commensal and animal isolates using both short- and long-read sequencing techniques. This enabled us to investigate the population plasmidome, core genome variation and genome architecture in detail, using a combination of machine learning, population genomics and genome-wide co-evolution analysis. We observed a high level of genome plasticity with large-scale inversions and heterogeneous chromosome sizes, collectively painting a high-resolution picture of the adaptive landscape of *E. faecium,* and identified plasmids as the main indicator for host-specificity. Given the increasing availability of long-read sequencing technologies, our approach could be widely applied to other human and animal pathogen populations to unravel fine-scale mechanisms of their evolution.

## Introduction

*Enterococcus faecium* is a Gram-positive bacterium that is ubiquitous as a commensal of the intestinal tract of humans and animals, as well as an opportunistic multi-drug resistant pathogen in debilitated patients (Van Tyne and Gilmore 2014). As commensal, *E. faecium* is a minority member of the intestinal microbiota mainly inhabiting the cecum and colon of adult humans (Tannock and Cook 2002). Although the functional role of *E. faecium* in the human intestinal tract is unclear, experiments in piglets suggest that enterococci may contribute to colonization resistance against pathogenic microorganisms (Willems and van Schaik 2009; Ness et al. 2014). In competition experiments *E. faecium* strains recovered from the intestinal microbiota of non-hospitalized persons outcompeted *E. faecium* strains from clinical infections in a murine intestinal colonization model (Montealegre et al. 2016). It was recently shown that part of this beneficial behavior resided in the production of secreted antigen A (SagA). SagA is a peptidoglycan hydrolase that generates muramyl-peptide fragments that were shown to protect the host (*Caenorhabditis elegans* and mice) against *Salmonella enterica* serotype Typhimurium and *Clostridium difficile* pathogenesis by upregulating host-barrier defenses (Rangan et al. 2016; Pedicord et al. 2016). As a pathogen *E. faecium* ranks among the most frequent causative agents of hospital-acquired infections, specifically central-line associated bloodstream infections (Weiner et al. 2016). The burden of disease due to *E. faecium* is augmented by the fact that *E. faecium* has acquired resistance against almost all available antibiotics, most notably against ampicillin, gentamicin and vancomycin, and, less frequently, against the more recently introduced antibiotics linezolid, daptomycin, and tigecycline (Guzman Prieto et al. 2016). This often results in multi-resistant phenotypes of isolates recovered from hospitalized patients, rendering infections with these clones difficult to treat. Antibiotic resistance, including vancomycin resistance, is not a feature exclusively found among hospitalized patient isolates with *E. faecium* isolates from farm animals also containing these resistance traits (Bonten et al. 2001). In addition to its multi-resistant nature, *E. faecium* is capable of colonizing patients to a high density, promoting frequent patient-to-patient transmission of clones (Ubeda et al. 2010; Ruiz-Garbajosa et al. 2012; Arias and Murray 2012). The full extent of *E. faecium* clonal dispersal has become apparent by whole genome sequencing (WGS)-based epidemiological studies. These have revealed frequent transmission of vancomycin-susceptible and vancomycin-resistant *E. faecium* among hospitalized patients and global dissemination of hospital-associated *E. faecium* clones (Raven et al. 2016; van Hal et al. 2016; Pinholt et al. 2015, 2017; Landerslev et al. 2016; Bender et al. 2016; Reuter et al. 2013; Brodrick et al. 2016; van Hal et al. 2017; Howden et al. 2013).

To obtain novel insights into the global population structure and recent evolution of *E. faecium* we developed novel genomic tools to analyze 1,684 *E. faecium* isolates from human (hospitalized patients, non-hospitalized persons) and animal (pet, farm and wild animals) sources. Our study thus develops a comprehensive approach which could be generally employed to study the detailed characteristics of the genomic landscape of a bacterial population. We explored the role of plasmids in the dynamics of adaptive evolution of *E. faecium* by analyzing the total plasmid content (plasmidome) of 62 isolates using a combination of short-read (Illumina) and long-read (Oxford Nanopore Technologies) sequencing. We employed a machine learning approach to predict the plasmidome of 1,582 *E. faecium* isolates (Arredondo-Alonso et al. 2018). We also characterized co-evolution of core genomic variation to identify signals of selection and adaptation. This revealed important genome-wide chromosomal rearrangements and showed that plasmid sequences, rather than variation in the *E. faecium* chromosome, are most informative for niche occupation in *E. faecium.*

## Results

### Core and accessory genome variability

Phylogenetic reconstruction of a set of 1,684 *E. faecium* isolates (Supplemental Table S1), based on a core genome alignment of 689,088 bp, confirmed the previous observations of a deep phylogenetic split that subdivided the *E. faecium* population into two clades, clade A (1,644 isolates) and clade B (40 isolates). The 40 Clade B isolates are highly diverse as they represent 32 different sequence types (STs) from a wide diversity of hosts (Supplemental Fig. S1, Supplemental Table S1). In order to maximize resolution in the core genomic analyses of the population, the 40 isolates belonging to clade B were removed and the remaining 1,644 isolates were studied in closer detail. The core genome alignment was filtered for recombination using BRATNextGen and the remaining variable sites were analyzed with hierBAPS to partition the population into Sequence Clusters (SCs) (Supplemental Table S1) (Croucher et al. 2013; Chewapreecha et al. 2014; von Mentzer et al. 2014). While core genes allowed the phylogenomic reconstruction of the *E. faecium* population structure (Fig. 1A, https://microreact.org/project/BJKGTJPTQ), they only covered a fraction of the full genome (978 orthologous groups). Analyses of accessory genes can provide a more compelling picture of the evolution of a species, for example by revealing niche-specific signals that represent the adaptation of different clones to similar environments (Richardson et al. 2018). The pan-genome of the 1,644 *E. faecium* isolates was composed of 30,855 orthologous groups and it was used to visualize isolate relatedness based on shared gene content analysis using PANINI (Fig. 1B, https://microreact.org/project/BJKGTJPTQ) (Abudahab et al. 2018). Isolates from hospitalized patients were largely separated from other sources. The distinct clustering of hospital isolates based on gene content in the pan-genome network is consistent with the core genome phylogeny. Despite the fact that hospital isolates form distinct clusters from isolates from other sources (the lower and left part of the network), they do not form a single or limited number of clusters in this network, indicating that the gene content of hospital isolates is highly diverse (Fig. 1B). The same is also true for pet isolates and isolates from non-hospitalized persons that are found throughout the upper part of the pan-genome network. This is also consistent with the distribution of these isolates among the core genome tree (Fig. 1A). In contrast, isolates from pigs and poultry in general cluster more closely together in their respective groups suggesting that gene content variation in these source groups is more limited.

**Figure 1:**
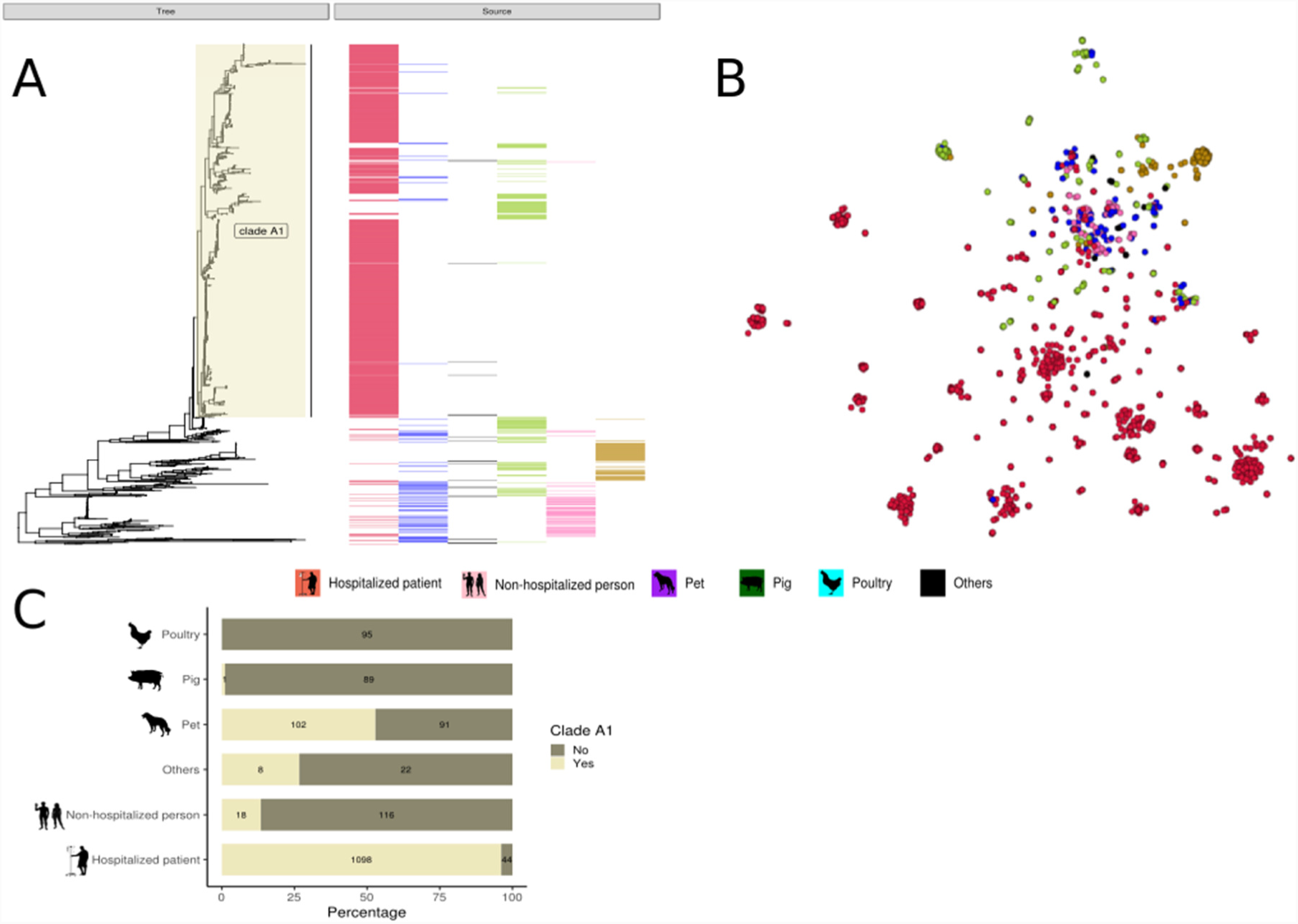
Population genetic analysis on 1,644 clade A *E. faecium* strains. A, RAxML tree based on 955 *E. faecium* core genes with the source distribution. B, PANINI network based on pan-genome colored according to sources. C, Percentage of sources in clade A1.

The distinct clustering of hospitalized patient isolates as described in the current study, has been widely reported and previous WGS analyses clustered hospital isolates in clade A1. In order to keep nomenclature harmonized with respect to the subpopulation of *E. faecium* hospital isolates, the clade (defined by the internal node 1227) that contained almost all (n = 1098; 96%) of the hospital isolates in this data set was designated clade A1 (Fig. 1A). Although hospital isolates also formed the largest proportion (1098/1227; 89%) of isolates in clade A1, a number of isolates (129/1227; 11%) from other sources, including isolates from non-hospitalized persons (n = 18), and pets (n = 102) was also present (Fig. 1C). The non-hospital source isolates in clade A1 were clearly dominated by isolates from dogs (78%), retrieved from different countries. The dogs from the Netherlands were randomly selected from an unbiased nationwide survey of healthy pet owners with no recent antibiotic usage history and who were themselves almost never colonized by hospital associated clones (van den Bunt et al. 2017).

### Chromosome and plasmidome sizes differ between hosts

To analyze the chromosome and plasmidome of the 1,644 clade A *E. faecium* isolates, we performed Oxford Nanopore sequencing and hybrid assembly on a selection of 62 *E. faecium* isolates. The 62 isolates were selected to capture the highest plasmidome variability present in our 1,644 clade A *E. faecium* isolates based on PlasmidSPAdes prediction (Antipov et al. 2016) and a homology search against a curated database of replication initiator proteins in enterococci (Clewell et al. 2014) (Supplemental Methods S3). We obtained 48 complete (finished) chromosomal sequences, and 305 plasmid and 6 phages in single circular contigs (Supplemental Table S2).

The 48 complete chromosomes ranged from 2.42 to 3.01 Mbp. Hospitalized patient isolates (n = 32) had the largest chromosomes (mean = 2.82 Mbp) whereas poultry isolates (n = 2) carried the smallest chromosomes (mean = 2.42 Mbp) in our set of complete genome sequences. Notably, hospital isolates may have up to 20% larger chromosome sizes than *E. faecium* from other sources, which highlights the considerable genomic flexibility of this organism.

The set of 305 complete plasmid sequences ranged in length from 1.93 to 293.85 kbp (median = 15.15 kbp, mean = 53.48 kbp) (Fig. 2A). Hospitalized patient isolates (n = 43) with complete plasmid sequences (n = 247) contained the highest number of plasmids (mean = 5.70) and their cumulative plasmid length was substantially larger compared to other isolation sources (mean = 308.01 kbp) (Fig. 2B and C).

**Figure 2:**
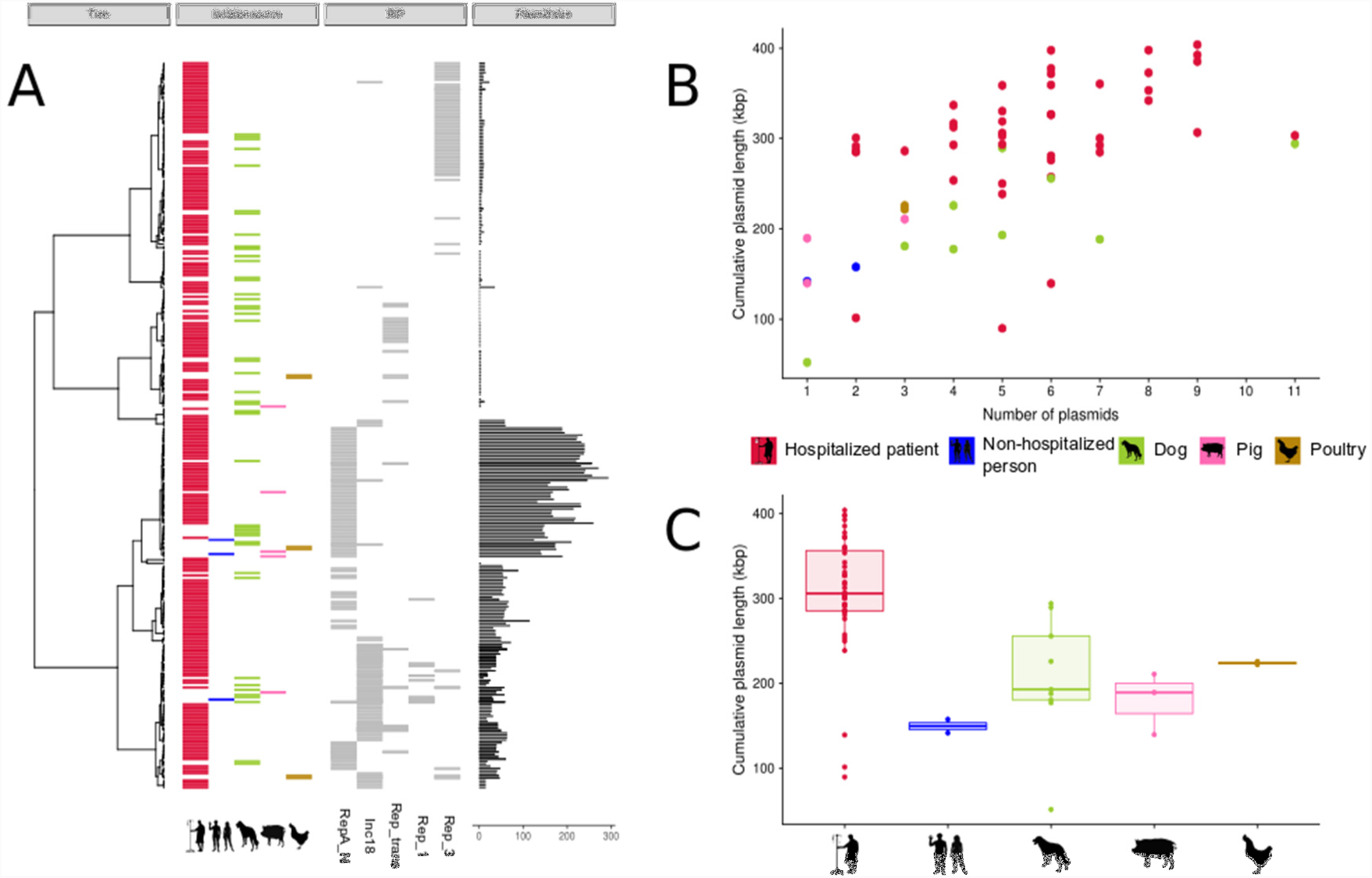
Genomic characterization of complete plasmid sequences (n = 305). A, Pairwise Mash distances (k = 21, s = 1,000) of the complete plasmid sequences (n = 305) were transformed into a distance matrix and clustered using hierarchical clustering (ward.D2). Node positions in the dendrogram were used to sort and represent in different panels the isolation source, replication initiator gene (RIP) and size (kbp) of the complete plasmid sequences. B, Number of plasmids (x-axis) and cumulative plasmid length (y-axis) are represented for 59 isolates with associated plasmid sequences. Hospitalized patients had the highest number of plasmids (n = 5.70) in contrast to non-hospitalized person (n = 1.5), pig (n = 1.67) and poultry (n = 3) but similar to the number of plasmids carried by dogs (n = 5.11). C, Boxplots of the average cumulative plasmid length (kbp) per isolation source.

### Genomic rearrangements in complete *E. faecium* genomes

Having fully assembled circular chromosomes in 48 isolates allowed us to assess the genomic organization in these isolates. This revealed four different large chromosomal rearrangements relative to the non-clade A1 strain E0139: i) type-1 (n = 1), inversion of 0.73 Mbp); ii) type-2 (n = 34), inversion of 1.2 Mbp; iii) type-3 (n = 2) two inversions of 0.38 Mbp; and type iv) (n = 1) inversion of 0.12 Mbp (Fig. 3A to D and Supplemental Fig. S2A to S2D).

**Figure 3:**
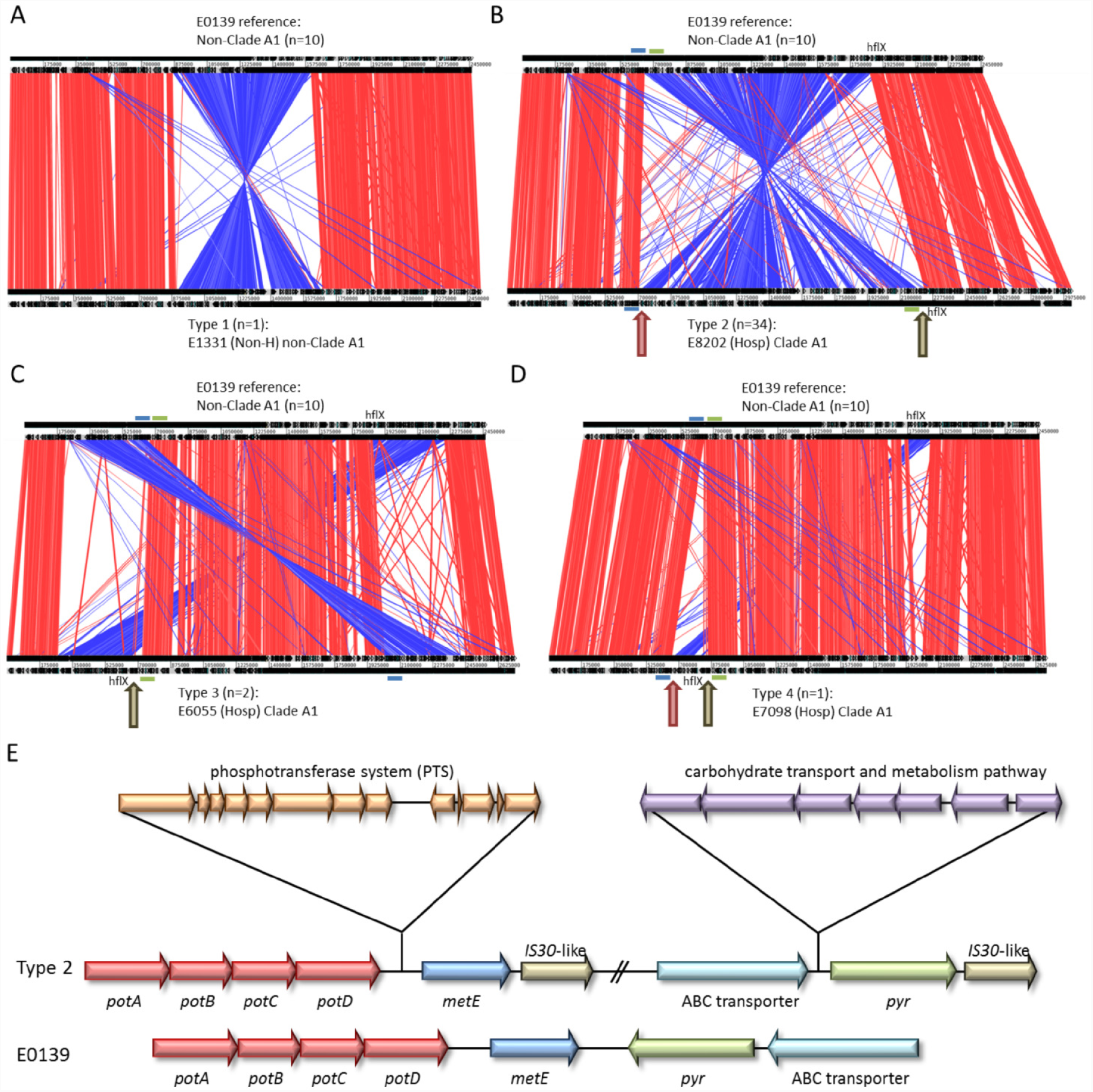
Analysis of genomic rearrangements using strain E0139 (non-clade A1) as a reference. A, type 1 genomic rearrangement (strain E1334). B, type 2 genomic rearrangement (strain E8202). C, type 3 genomic rearrangement (strain E6055). D, type 4 genomic rearrangement (strain E7098). E, Schematic representation of insertion and inversion sites in the most predominant type 2 genomic rearrangement (strain E8202) compared to reference strain E0139. Blue bar: Pot operon; Green bar: ABC transporter; red arrow: insertion site phosphotransferase system encoding genomic island (Zhang et al. 2013b); brown arrow: insertion site carbohydrate transport system encoding genomic island (Heikens et al. 2008).

Except for the type-1 inversion, all these genomic rearrangements were exclusively observed in clade A1 isolates (Supplemental Fig. S3), and may be associated with adaptation of *E. faecium* to nosocomial ecology. In clade A1 isolates we detected two previously described genomic islands inserted adjacent to the genomic rearrangements (Heikens et al. 2008; Zhang et al. 2013b) (Fig. 3E). Both genomic islands have been described to be enriched among clade A1 isolates. One genomic island, putatively encoding a carbohydrate transport and metabolism pathway (Heikens et al. 2008), was always found to be located downstream of an ABC transporter, while the other element encoding a phosphate transport (PTS) system (Zhang et al. 2013b) was always found to be located downstream of a polyamine transport system (Fig. 3E). In all cases, the genomic rearrangement was flanked by an *IS30*-like transposon downstream of a methionine synthase (*metE*) and a pyridine nucleotide-disulfide oxidoreductase (*pyr*) (Fig. 3E). We determined the presence of the two genomic islands with insertion sites in the draft genomes of the remaining 1,596 isolates and confirmed that the genomic islands were always inserted at the same position in 93% of the clade A1 isolates. In addition, the insertions were also identified in 57 (14%) non-clade A1 isolates, which mainly represented dog isolates (n = 43) (Supplemental Fig. S3). In contrast, a branch in clade A1 consisting of 68 isolates, of which 55 originated from dogs, lacked both genomic island insertions (Supplemental Fig. S3). In isolates that lacked insertions of the two genomic islands the genomes were organized in a manner comparable to the non-clade A1 isolate E0139, i.e. without inversion (Supplemental Fig. S3).

### Plasmids exhibit extensive mosaicism

The 305 completely sequenced plasmids were first characterized with a standard (Clewell et al. 2014) categorization based on the: i) presence of replication initiator proteins (RIP) (Supplemental Table S3), and ii) presence of relaxases (MOB) (Supplemental Table S4).

A considerable proportion of plasmids (48/294, 16%) were multireplicon plasmids, with plasmids encoding up to four different RIP gene families. This was most prominent in Rep_1 and Inc18 family plasmids, which contained at least one other RIP with a frequency of 1.0 (8/8) and 0.53 (30/57) (Supplemental Fig. S4A), respectively. The predominant RIP family RepA_N (n = 82) was mainly encoded on large plasmids (mean plasmid length = 155.3 kbp) and was less frequently associated with other RIP sequences (n = 15, 18%) (Supplemental Fig. S4A). Plasmids encoding the Rep_3 family (n = 56, mean plasmid length = 12.4 kbp) and Rep_trans (n = 24, mean plasmid length = 25.7 kbp) were less frequently present in multireplicon plasmids (n = 6, 11%) (Supplemental Fig. S4A). No RIP family could be characterized for 11 plasmids (mean plasmid length = 9.6 kbp).

The extreme modularity of *E. faecium* plasmids became even more apparent when relaxase gene families were linked to the fully sequenced plasmids. All identified relaxases co-occurred in plasmids with different RIP genes and even in multireplicon plasmids (Supplemental Fig. S4). In total we observed 46 different Rep-relaxase combinations (Supplemental Fig. S4A). A more extensive characterization of the mosaicism of plasmid sequences is available in the Supplementary Results S1.

### The plasmidome of *E. faecium* differs substantially between hosts

To predict the plasmidome content of our entire sequenced population, we used mlplasmids which provides a support-vector machine classifier to label contigs as plasmid or chromosome derived (Arredondo-Alonso et al. 2018). Mlplasmids predicted 165,200 and 68,363 contigs as having a chromosome- and plasmid-origin, respectively (Supplementary Table S5). The average number of base-pairs predicted as plasmid- (240,324 bp ∼ 52 contigs) or chromosome-derived (2,619,359 bp ∼ 113 contigs) was in line with the expected genome size of *E. faecium*. Mlplasmids did not predict plasmid-derived contigs in four isolates, from four different isolation sources (non-hospitalized, dog, hospital patient and sea cucumber), including one known plasmid-free isolate 64/3 (in this study named E2364) (Bender et al. 2015).

We observed significant differences in the number of base-pairs predicted as plasmid- and chromosome-derived depending on the *E. faecium* host (Kruskall-Wallis test, *P* < 0.05) (Fig. 4A and 4B). Predicted chromosome and plasmidome size of isolates from hospitalized patients were considerably larger (mean = 2.68 Mbp and 276.16 kbp) compared to other isolation sources (Fig. 4A and 4B). This finding is in line with previous reports which showed that isolates from clade A1 are enriched for mobile genetic elements (Lebreton et al. 2013; Buultjens et al. 2017).

**Figure 4:**
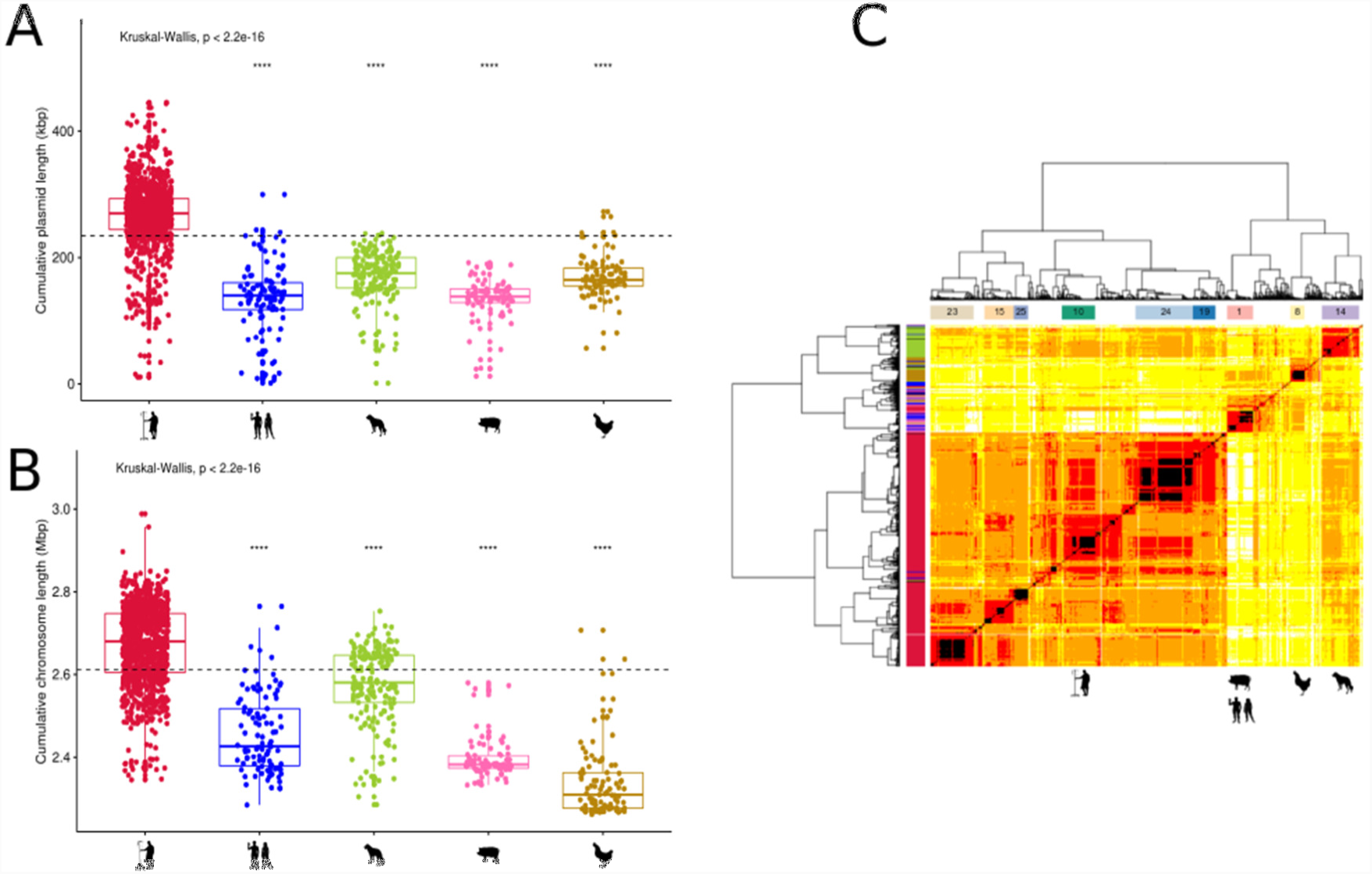
Predicted plasmidome of 1,644 *E. faecium* isolates. A, Boxplot of the number of base-pairs (kbp) predicted as plasmid-derived per isolation source. Horizontal dashed line indicates the mean cumulative plasmid length across all the groups. B, Boxplot of the number of base-pairs (Mbp) predicted as chromosome-derived per isolation source. Horizontal dashed line indicates the mean cumulative chromosome length across all the groups. C) Pairwise Mash distances (k = 21, s = 1,000) of plasmid-predicted contigs in 1,607 isolates were transformed into a distance matrix and clustered using hierarchical clustering (ward.D2). Based on the quantile function of our defined Gamma distribution, we grouped isolates in five blocks: black (0-0.01), red (0.01-0.25), orange(0.25-0.5), yellow (0.5-0.75) and white (0.75-1.0). Dissimilarity matrix of the isolates was visualized by a heatmap coloured based on the previous blocks. We incorporated the defined plasmid populations (n = 9) and isolation source information on top and left dendrograms respectively.

In order to assess the extent of association between plasmid content and isolation source, we computed pairwise distances of isolates based on the k-mer content of their predicted plasmidome, and computed a neighbor-joining tree (bioNJ) (Fig. 5A and Supplemental Fig. S5) (see Material and Methods). In this analysis 37 isolates were excluded as they showed no signs of plasmid carriage signatures based on their distribution of pairwise distances. To estimate clusters of isolates with a similar plasmidome content, we used hierarchical clustering (ward.D2) to group isolates based on pairwise distances and divided the resulting dendrogram into distinct groups with an optimal number of 26 clusters (avg. silhouette width = 0.45) (Supplemental Fig. S6A). From these clusters, we selected plasmid populations (n = 9) that fulfilled two criteria: i) cluster size (larger than 50 isolates) and ii) goodness-of-fit measure (avg. silhouette width higher than 0.3) (Supplemental Fig. S6B).

**Figure 5:**
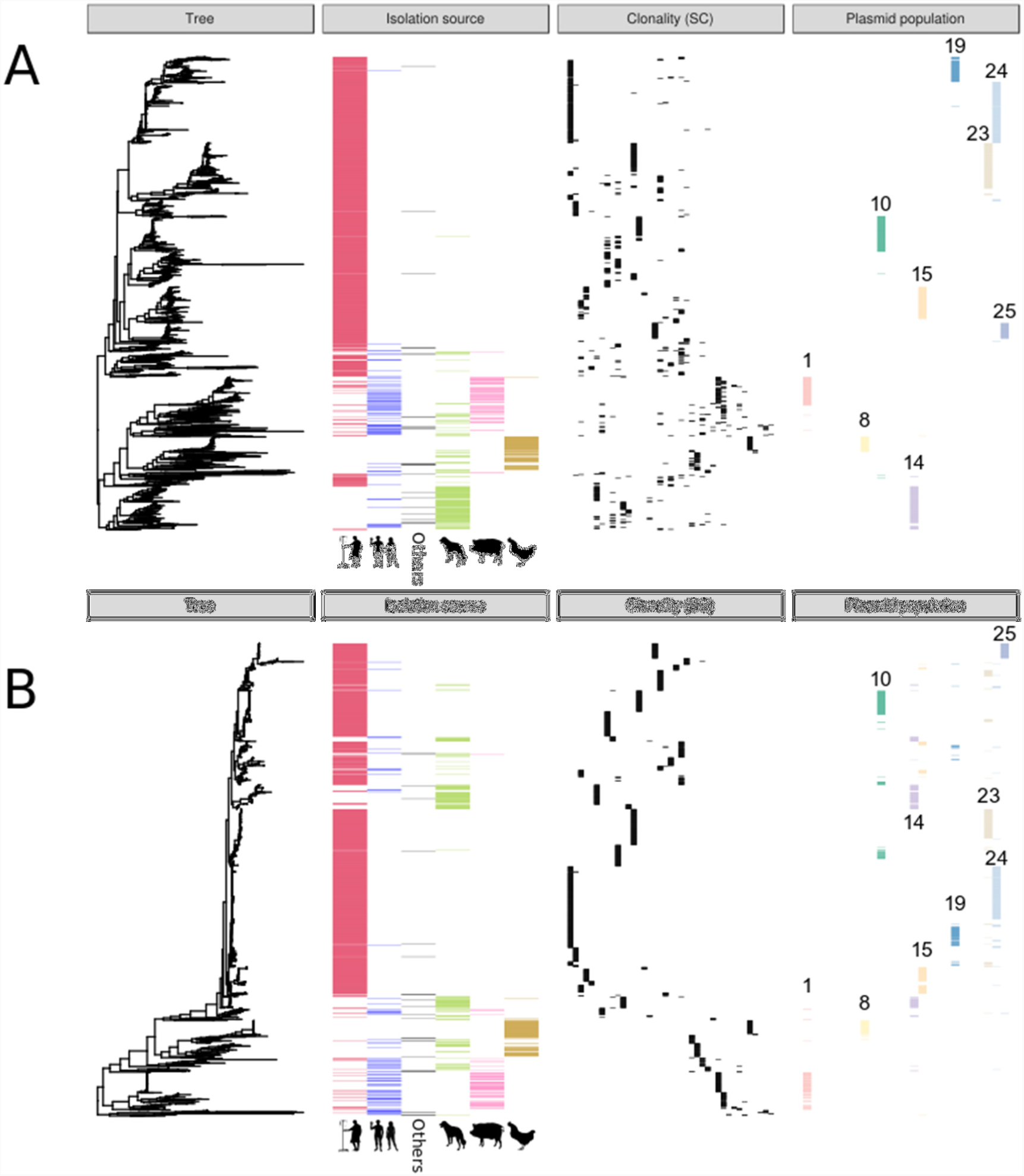
Comparison of reconstructed *E. faecium* phylogeny based on plasmid- and core-genome sequences. We incorporated three different panels of information: isolation source, clonality (SC) and plasmid population. Each of the horizontal bars in the panel corresponds to a different group (isolation source, SC or plasmid population). A, bioNJ tree based on the dissimilarity matrix of Mash distances (k = 21, n = 1,000) of 1,607 isolates uniquely considering plasmid-predicted contigs. B, ML core-genome tree based on a core genome alignment of 689,088 nts.

Plasmid subpopulation 1 (size = 99, avg. sil. width = 0.75) was mainly populated by isolates from pigs and non-hospitalized persons (Bonferroni corrected *P* < 0.05) (Fig. 4C and Supplemental Fig. S6B) with heterogeneous SCs (Simpson index = 0.53) suggesting horizontal transmission of plasmid sequences between pig and non-hospitalized persons (Supplemental Fig. S6C). Chicken isolates were overrepresented in plasmid population 8 (size = 52, avg. silhouette width = 0.82) (Bonferroni corrected *P* < 0.05) (Fig. 4C and Supplemental Fig. S6B) but SC homogeneity (Simpson index = 0.21) suggested that dissemination of this plasmid population was mainly driven by clonal transmission (Supplemental Fig. S6C). Dog isolates were overrepresented in plasmid population 14 (size = 139, avg. silhouette width = 0.41) (Fig. 4C and Supplemental Fig. S6B) with heterogeneous SCs indicating horizontal transmission of plasmid sequences (Simpson index = 0.73) (Supplemental Fig. S6C). Hospitalized patient isolates were overrepresented in six different plasmid subpopulations (Fig. 4C and Supplemental Fig. S6B). Based on SC diversity, we observed that dissemination of plasmid populations 19, 24 and 25 (size = 83, 214 and 53; avg. silhouette width = 0.31, 0.64 and 0.90; Simpson index = 0.30, 0.15 and 0.04 respectively) were mainly driven by clonal transmission (Supplemental Fig. S6C and Supplemental Results S2). However, plasmid population 10, 15 and 23 exhibited higher SC diversity (size = 125, 110 and 160; avg. silhouette width = 0.53, 0.60 and 0.64; Simpson index = 0.60, 0.73 and 0.56 respectively), suggesting horizontal transmission events of plasmid sequences between isolates from different SCs (Supplemental Fig. S6C and Supplemental Results S2).

We further investigated the influence of plasmids on the *E. faecium* phylogeny by transforming Mash distances into a bioNJ tree for three different input data: i) plasmid-predicted contigs (see above), ii) chromosome-predicted contigs and iii) whole-genome assembly. Reconstructed phylogenies were compared against the core genome tree. We observed that phylogeny inferred by plasmid-derived contigs showed the highest Kendall and Colijn (KC) distance when considering either only tree topology (KC distance = 12,206.81) or a balanced measure between tree topology and branch length (KC distance = 6,103.53). This is highly suggestive of major differences in the evolutionary trajectories of the core genome and plasmid sequences of *E. faecium* (Supplemental Fig. S7).

To assess the extent of association between plasmid content and isolation source, we explored the orthologous groups present in the previously described isolates (n = 1,607). The total plasmid coding capacity (pan-plasmidome) amounted to 5,458 orthologous genes. Only a minor fraction of genes (n = 454) were found in more than 15% of the isolates. We further investigated functional differences in plasmid orthologous groups between *E. faecium* hosts with a gene enrichment analysis using Scoary (Brynildsrud et al. 2016). Plasmidomes of hospitalized patient isolates were enriched (Specificity & Sensitivity > 80%) for a large pool of genes (n = 589, OR > 1) (Supplementary Fig. S8) with a variety of functions, including a bacteriocin-like protein (*bacA)* previously described as plasmid-related in *E. faecalis* (Kurushima et al. 2016) and antibiotic-resistance related genes such as *aacA-aphD* or efflux permease ABC transporters (Supplementary Table S6). Poultry and pig isolates showed a pool of 155 and 100 enriched plasmid genes (OR > 1; Specificity & Sensitivity > 80%), respectively (Supplementary Fig. S8). Pig-associated plasmid genes included genes involved in copper resistance (*tcrB, tcrY, tcrZ, cueO*) which was a growth-promoting agent commonly used in piglets (Hasman et al. 2006). In poultry two genes encoding a choloylglycine-hydrolase protein, potentially involved in the tolerance of high bile salt concentrations and a cold-shock protein (*cspC)* were enriched. A detailed characterization of plasmid enriched genes is available in the Supplementary Results S3.

We further explored and quantified the association between isolation source and genomic component by comparing the distribution of pairwise distances of: whole-genome contigs, chromosome-derived contigs and plasmid-derived contigs. We hypothesized that pairwise distances between isolates belonging to the same isolation source are lower than pairwise distances from isolates of different sources and that the difference in magnitude reflects how informative each component (chromosome, plasmid, whole-genome) is for host-specificity. We computed the average pairwise distance of the following combinations: i) pairs of isolates from a same isolation source (within-host group), ii) pairs of isolates belonging to different isolation sources (between-host group) and iii) pairs of isolates randomly selected (random group)). We repeated this process following a bootstrap approach (100 iterations) and assessed whether the distribution of within-host and between-host groups pairwise distances statistically differed (one-way anova test) from the random group (Fig. 6 and Supplemental Fig. S9).

**Figure 6:**
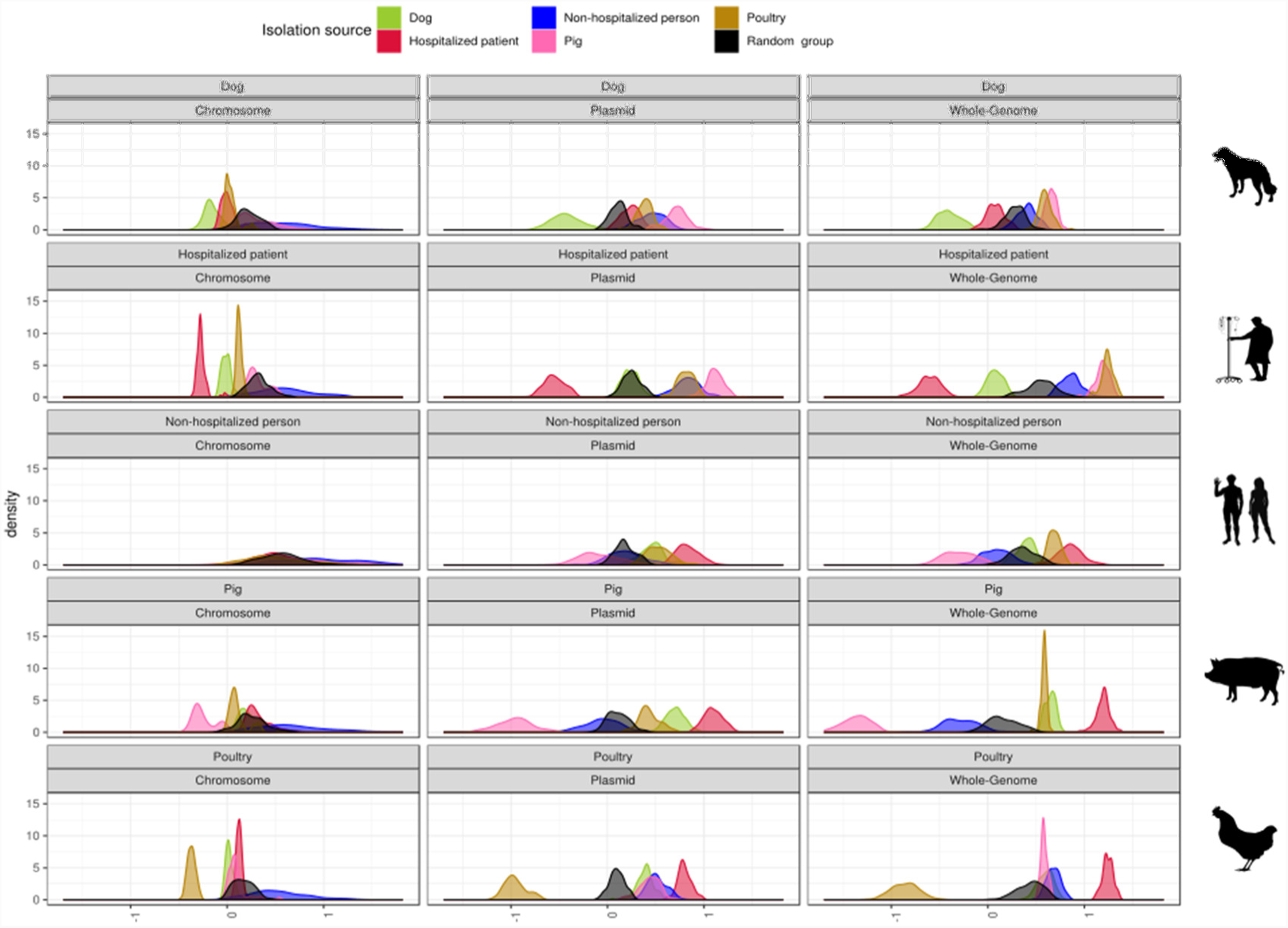
Distribution of average pairwise distances based on isolation source and genomic component. Pairwise Mash distances from whole-genome contigs, plasmid-predicted contigs, chromosome-predicted contigs were scaled and compared between each isolation source. We used a bootstrap approach (100 iterations), to compute the average pairwise distance of: i) pairs of isolates belonging to the same host, ii) pairs of isolates belonging to different hosts and iii) pairs of isolates belonging to random isolates.

We observed that the whole-genome contigs explained most of the host specificity in all the isolation sources, with the exception of non-hospitalized persons, as shown by the highest observed difference in mean pairwise distance between isolates belonging to the same isolation source and randomly selected sources (within-host pairwise distances) (Fig. 6 and Supplemental Fig. S9). Except for non-hospitalized persons, the separation into plasmid- and chromosome-derived contigs revealed that the contribution of plasmids was higher than the chromosome-component for all isolation sources, as exemplified by a higher observed mean difference in within-host and between-host groups (Fig. 6 and Supplemental Fig. S9). Furthermore, we observed that in some cases the plasmid component showed even higher levels of dissimilarity than the whole-genome component when comparing between-host groups (Supplemental Fig. S9). Most notably, we observed significant similarities in the whole genome (diff. mean = 0.20, *P* < 0.05) and chromosome (diff. mean = 0.21, *P* < 0.05) between dog and hospitalized patient isolates but a significant dissimilarity of the plasmid sequences between these host groups (diff. mean = −0.13, *P* < 0.05). Host specificity determined by plasmid sequences was highest in pig isolates (diff. mean = 1.07, *P* < 0.05) and poultry isolates (diff. mean = 1.05, *P* < 0.05) (Supplemental Fig. S9). Furthermore, we observed significant differences in the average plasmid pairwise distance of pig and non-hospitalized isolates (diff. mean = 0.15, *P* < 0.05) suggesting the exchange of plasmid sequences between these two hosts (Supplemental Fig. S9).

### Co-evolution of core genomic variation

Following the analysis of the population structure, genome synteny and the plasmidome, we next aimed to identify signals of selection acting to shape co-evolution of SNPs. Co-evolved SNPs can facilitate adaptation to different environments when sequentially selected mutations in adaptive elements decrease the actual fitness costs of individual mutations (antagonistic epistasis) and thus have a beneficial effect on fitness (Lagator et al. 2014).

We performed genome-wide analysis of SNP co-evolution using Direct Coupling Analysis (SuperDCA) on the core genome alignment of *E. faecium* to identify likely candidates of sequentially selected or coupled mutations not in strong linkage disequilibrium due to chromosomal proximity (Puranen et al. 2018). SuperDCA was applied to a core-genome alignment generated using the Harvest suite, including the 1,644 Clade A isolates and the complete *E. faecium* AUS0004 (accession number CP003351) as reference (Lam et al. 2012). All locus tags mentioned below refer to AUS0004.

### Analysis of ten loci with the highest number of linked SNPs

SuperDCA identified 262,877 significant couplings between SNPs with a distance of > 1 kbp (Supplemental Table S7). Ten loci with the highest number of linked SNPs were plotted individually to obtain insight in the distribution over the AUS004 core genome (Supplemental Fig. S10). This showed that loci 3 and 10 (representing genes *EFAU004_02173* and *EFAU004_02178*, respectively), which were located in close proximity to each other, had a similar pattern of coupled SNPs with genes *EFAU004_00665* to *EFAU004_00675* located at a large chromosomal distance of around 500 kbp in the AUS0004 core genome. In total, 223 SNPs in genes *EFAU004_02173* to *EFAU004_02178*, referred to as region-1, were linked to genes *EFAU004_00665* to *EFAU004_00675*, referred to as region-2, (Fig. 7A and Supplemental Table S8).

**Figure 7:**
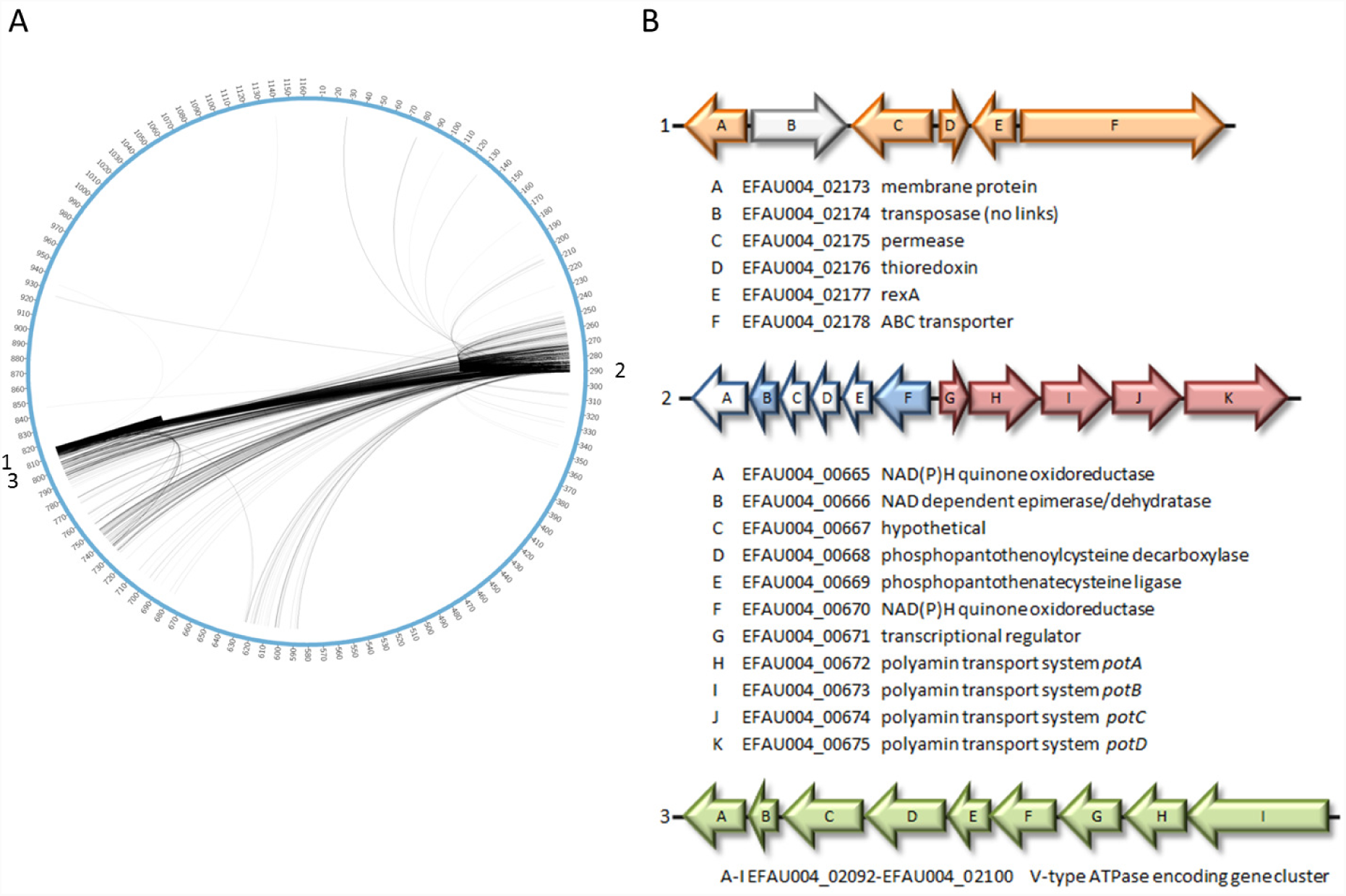
Distribution and genomic organization of coupled SNPs. A, All 17,236 linked SNPs of the genes contained in region 1 and 2. B, Genomic organization and annotation of genes from region 1-3 in AUS0004.

Careful examination of the exact localization of the genes in region-1 and region-2 revealed that these regions were part of the large chromosomal rearrangement described above. This means that in the majority of non-clade A1 isolates region-1 and region-2 are located adjacent to each other as schematically indicated for *E. faecium* strain E0139, isolated from a non-hospitalized person (Fig. 3E).

Region-1 is predicted to encode a thioredoxin (Trx) (*EFAU004_02176*) and the transcriptional regulator Rex1 (*EFAU004_02177*). Trx and Rex1 are likely involved in redox homeostasis of the bacterial cell, which was found to be critical for DNA synthesis and defense against oxidative stress (Wang et al. 2008) (Fig. 7B). The genes from region-2 can be split into two distinct gene clusters based on their putative biological functions (Fig. 7B). Genes *EFAU004_00665* to *EFAU004_00670* include genes which are predicted to encode a NAD(P)H:quinone oxidoreductase involved in respiration and coenzyme A biosynthesis. Genes *EFAU004_00671* to *EFAU004_00675* seems to be organized in an operon and display similarity with a polyamine transport system. In *Streptococcus pneumoniae* polyamines are pivotal in survival strategies in the host when bacteria are confronted with stress conditions like temperature shock, oxidative stress, or choline limitation (Shah et al. 2008).

All genes from region-2 are also predicted to have co-evolved with a gene cluster close to region-1, indicated as region-3 (Fig. 7). This region-3 is predicted to encode a membrane bound V-type ATPase. This V-type ATPase has been studied in detail in *Enterococcus hirae* (Murata et al. 2001). The V-type ATPase belong to the family of proton pumps and are involved in the translocation of Na^*+*^ or H^*+*^ over the cell membrane by using the energy of ATP. In *E. hirae* the V-type ATPase appeared highly expressed under stress conditions like high pH and plays an important role in sodium homeostasis under these conditions. A more detailed description is available in the Supplementary Results S4.

## Discussion

*E. faecium* is widely distributed among humans, animals and the environment. It is thought that the maintenance of microbial communities facilitating exchange of plasmids, phages and conjugative transposons has allowed them to rapidly adapt to and increase their fitness in changing environmental conditions (Guzman Prieto et al. 2016). In this study, we used a combination of ONT and Illumina short-read technologies to paint a more comprehensive picture of how the nosocomial pathogen *E. faecium* has evolved in different host environments and how interlinked its plasmid population is.

We investigated the role of plasmids in host specificity by predicting the plasmidome content of our *E. faecium* collection using a machine-learning classifier specifically designed at a species-level (Arredondo-Alonso et al. 2018). We observed that total plasmid and chromosome size of isolates from hospitalized patients was substantially larger compared to animal isolates and isolates from non-hospitalized persons. Clustering of pairwise Mash distances of predicted plasmid sequences revealed a high-level of diversity and modularity in *E. faecium* plasmids. We then estimated the potential contribution of different genomic components (whole-genome, chromosome, and plasmid) to host specificity. We observed that the plasmid component drives host specificity in dogs and hospitalized patients while their whole genome and chromosome share a common evolutionary history. In line with previous reports (Willems et al. 1999; Freitas et al. 2011), we observed that non-hospitalized patient isolates shared a plasmid population with pig isolates, which indicates exchange of plasmids between both hosts. Host-specificity driven by plasmid sequences was highest in pigs and poultry isolates and significantly differed from other hosts. Hospitalized patient isolates were associated with six different plasmid populations indicating the presence of different plasmid configurations within the hospital environment. Furthermore, the plasmidome of clinical isolates are highly dissimilar to those from other hosts. These data indicate that the plasmid component of the *E. faecium* pan-genome plays a major role in adaptation and the emergence of this organism as a nosocomial pathogen of major importance.

Exploration of plasmid genes specifically present in each of the isolation sources revealed the presence of a widely-spread bacteriocin-like gene (*bacA*) in our set of hospitalized isolates that was rarely present in other isolation sources. As previously reported for *E. faecalis*, *bacA* can act as a sophisticated toxin-antitoxin system, in which not only daughter but also co-existing *bacA* plasmid-free cells are excluded from the population (Kurushima et al. 2016). As such, this *bacA* gene may have contributed to the ecological dominance of the distinct subpopulation of *E. faecium* hospital isolates in patients. Linkage of specific plasmid genes to particular hosts was not restricted to hospitalized patients. We also found a set of copper resistance genes enriched in *E. faecium* plasmids from pig isolates. Since copper was used as a growth-promoter agent in piglets and high levels of copper are toxic for most bacterial species, the acquisition of copper resistance genes may have contributed to the adaptation of *E. faecium* to environmental constraints imposed by pig farming. Recently, Gouliouris et al. 2018 also described the same copper-resistance operon to be overrepresented in pig isolates and thus confirming that this set of plasmid-encoding genes have played an important role in *E. faecium* survival in farms (Gouliouris et al. 2018). We also identified a choloylglycine-hydrolase gene widely present in poultry isolates. *E. faecium* has been previously characterized as one of the microorganisms with highest levels of bile salt hydrolase activity in the intestine of chickens (Knarreborg et al. 2002) and capable of developing new mechanisms to tolerate high concentration of bile salts (Zhang et al. 2013a). The choloylglycine-hydrolase gene described here could be functionally responsible for bile tolerance of poultry isolates. We also observed a plasmid gene (*cspC)* involved in cold-shock response and NaCl, pH, and ethanol stress in Clostridium (Derman et al. 2015). The observation that plasmid sequences are highly informative for host-specificity and that particular plasmid genes may have a clear benefit for *E. faecium* in particular hosts suggest that the distribution of plasmid genes among *E. faecium* isolates is regulated by complex ecological constraints, thus have contributed to host adaptation, rather than by opportunity arising from physical interaction between different host types.

The population plasmidome analysis further disclosed the high modularity of *E. faecium* plasmids. The high number of multireplicon plasmids consisting of several combinations of Rip families confirmed the high levels of mosaicism also previously observed for *E. faecium* plasmids which makes it infeasible to classify *Enterococcal* plasmids based on Rip-schemes (Freitas et al. 2016). More importantly, it illustrates the heterogeneity and flexibility of these extra-chromosomal elements in which genetic modules are frequently swapped.

*E. faecium* hospital isolates do not only differ in plasmid content but, as reported before, they also have a distinct evolutionary history that resulted in the evolution of the distinct *E. faecium* subpopulation, referred to as clade A1 (Lebreton et al. 2013). Other than isolates from hospitalized patients, only isolates from pets (mainly dogs) are also frequently associated with this clade, which is consistent with previous studies (de Regt et al. 2012; van den Bunt et al. 2017). Approximately half of the dog isolates fall into the hospital associated clade A1 and this is unlikely to originate from epidemiological links between the dogs in the main dog SC since they originate from three different countries (Netherlands, Denmark and the UK) and the Dutch dog isolates stem from a nationwide unbiased survey (van den Bunt et al. 2017). Furthermore, based on the PANINI network and plasmid pairwise distance measurements, the accessory genome of dogs and patients isolates differ. This identifies dogs as a possible source from which hospital-adapted *E. faecium* clones have evolved but which subsequently acquired additional mobile genetic elements that contributed to their success as nosocomial pathogens.

Human community isolates from non-hospitalized patients are widely dispersed over the phylogenetic tree outside clade A1. In previous reports, isolates from non-hospitalized persons dominated clade B (Lebreton et al. 2013) and were supplemented by isolates from other sources (Raven et al. 2016; Gouliouris et al. 2018). In the current study, most isolates from non-hospitalized persons cluster outside clade B and cluster near isolates from a variety of different sources, which may indicate either frequent exchange or easy adaptation to different ecological niches. This suggests that human community isolates are a result of colonization by *E. faecium* clones from different sources (poultry, pet, pigs), and that this may have occurred relatively recently, and or is still ongoing, preventing (sub)speciation events that lead to the evolution of human-specific (non-hospital associated) clones.

Farm animal isolates, represented in this study mostly by isolates from poultry and pigs, clustered in clade A distinct from the hospital clade A1 in polyphyletic groups suggesting that there is no single farm animal associated clade A2 as reported previously (Lebreton et al. 2013). A core genome analysis by Raven et al. similarly did not find support for the presumed subdivision of clade A into clade A1 and A2 (Raven et al. 2016). Pig and poultry isolates were grouped in a limited number of distinct SCs, with 88% of pig isolates grouping in SC 29 and 30 and 93% of poultry isolates grouping in SC24, 25 and 35 (Supplemental Table S1). This clearly indicates host-specific evolution of *E. faecium* in distinct ecotypes, something that was suggested decades ago based on DNA fingerprint data (Willems et al. 2000) and later confirmed using MLST (Willems et al. 2012) and whole-genome sequencing (Gouliouris et al. 2018). If we consider SCs 29 and 30 as genuine pig-associated lineages then the isolates from patients (n=11) and non-hospitalized persons (n=33) in these SC groups may represent spillover events from these farm animals.

Pangenomic analysis showed that the hospital associated *E. faecium* population harbors a very diverse accessory genome, compared with any other source, including animals. Hospital associated *E. faecium* clones seem to be unable to successfully establish themselves outside hospitals. A parsimonious interpretation of these two observations is that the hospital associated *E. faecium* population continuously imports novel genes as the isolates circulate and continues to form local clouds of diversity by localized transmission. This further suggests that there is an ample opportunity for new hospital associated *E. faecium* clones to emerge in the future, once the necessary fitness elements come into place.

Traces of adaptations are not only displayed by specific signatures of the population plasmidome or pangenome but also by the appearance of co-evolved SNPs in three chromosomal regions; region-1, region-2 and region-3. One of the main findings of this analysis was that co-evolved SNPs in these chromosomal regions were located on the borders of a large genomic rearrangement and therefore adjacent to each other in the majority of non-clade A1 isolates but at a large distance in clade A1 isolates. In all cases, the genomic rearrangement is flanked by an *IS30*-like transposon downstream *metE* and upstream *pyr*. The predominant chromosomal rearrangement, type-2, identified in the current study is also present in other publicly available completed genomes such as *E. faecium* DO (NC_017960.1) (Qin et al. 2012), *E. faecium* Aus0085 (NC_021994.1) (Lam et al. 2013) or *E. faecium* E39 (NZ_CP011281.1). For the isolates in our study for which we only have a draft genome, it is difficult to determine whether genomic rearrangements have occurred. However, in isolates in which an *IS30*-like element was identified there was always a contig break with this element at the border downstream the *metE* or *pyr* gene suggesting a genomic rearrangement. A chromosomal rearrangement in *E. faecium* was observed in the first completely sequenced strain AUS0004, where it was hypothesized that it occurred between 2 phage elements resulting in a replichore imbalance (Lam et al. 2012). However, we did not observe a replichore imbalance in any of the 38 completely assembled genomes with large chromosomal rearrangements.

The gene clusters located in the chromosomal regions-1, -2, and -3 that harbored co-evolved SNPs clearly encode common functions. The thioredoxin pathway, Rex1 (region-1) and the NAD(P)H dependent oxidoreductase/epimerase proteins (region-2) are all dependent on the oxidation/reduction of NAD^*+*^/NADH and involved in redox homeostasis, while the polyamine transport system (region-2) and V-type ATPase (region-3) are both involved in ATP hydrolysis/synthesis. The fact that in non-clade A1 isolates chromosomal regions 1, 2 and 3 are located adjacent to each other explains the high number of statistically significantly linked SNPs between the genes in these two regions, as they likely have co-evolved in fairly tight LD. Conversely, this finding illustrates that the co-evolutionary analysis of SNPs can uncover inversions in the chromosome, which can play a role in adaptation to new environments by changing the gene expression landscape and more particularly to stress conditions, like the hospital environment where bacteria are exposed to different stressors including high concentrations of antibiotics, detergents and antiseptics. To our knowledge, identification of genomic rearrangements by coupling analysis of SNPs has not been previously discussed in the literature, and the expected rapid increase in the availability of long-read sequences will facilitate development of new methods to identify candidates of inversions associated with selective advantage.

Taken together, we propose that *E. faecium* specialization into clades stems from genetic adaptivity promoted by high-level gene flux through acquisition and loss of plasmids carrying adaptive elements as well as evolutionary volatile genomic rearrangements. This means that the genomic signature of *E. faecium* isolates is highly dynamic with a continuous flux of accessory genes and horizontal gene transfer events. This genome plasticity and evolutionary flexibility of *E. faecium* is not only visible in accessory genome dynamics but also in the core genome where interactions between polymorphisms likely play a role in continuously fine-tuning the genetic configuration thereby facilitating adaptations to changing conditions. Combining extensive short- and long-read sequencing of a large collection of isolates from a diverse set of hosts, as reported here for *E. faecium*, may serve as a broadly applicable approach to study the effects of adaptation and selection on optimizing the genome architecture in successful lineages in evolutionary and ecologically diverse populations.

## Material and Methods

### Genomic DNA Sequencing and assembly

Detailed description of Illumina and ONT sequencing is available at Supplementary Methods S1-S5 and in Arredondo-Alonso et al. (Arredondo-Alonso et al. 2018) which includes a full description on ONT selection of *E. faecium* isolates (n = 62) and consecutive hybrid assembly using Unicycler (Wick et al. 2017).

### Population genomic analysis

Pangenomes for the entire genome dataset (1,684 strains) and the clade A dataset (1,644 strains) were created using Roary (Page et al. 2015) with default settings. A core gene alignment was generated using the –mafft option in Roary resulting in a core gene alignment of 859 genes for the entire dataset and of 978 genes for the clade A dataset. PANINI was run with the default settings using the Roary gene presence/absence matrix to generate a network of isolates based on the similarity of their gene content (Abudahab et al. 2018). To estimate recombination events and to remove them from the core genome alignment we used BratNextGen with default settings including 20 HMM iterations, 100 permutations run in parallel on a cluster, and 5% significance level similar to earlier publications (Marttinen et al. 2012; de Been et al. 2013). To determine sequence clusters in the core genome alignment where significant recombinations had been removed, we used 5 estimation runs of the hierBAPS method (Cheng et al. 2013) with 3 levels of hierarchy and the prior upper bound for the number of clusters ranging in the interval [50-200]. All runs converged to the same estimate of the posterior mode clustering. To estimate a phylogenetic tree we used RAxML (Stamatakis 2014) with the GTR+Gamma model on core gene alignment stripped of recombination. Bootstrap option was disabled in RAxML due to an extremely long runtime.

### *E. faecium* genome organization

To unravel rearrangements in the chromosome, we considered a clade-A2 isolate from a non-hospitalized person (E0139) with a complete chromosome sequence as reference. We only considered isolates with a complete circular chromosome (n = 48). Pairwise chromosomal comparisons were computed using blastn (version 2.7.1+) and alignments were visualized using the Artemis Comparison tool (version 17.0.1). The insertions of two genomic islands encoding a phosphotransferase system and carbohydrate transport system and their putative co-localization with the pot operon and ABC transporter, respectively was determined by BLAST. Additionally, for each chromosomal rearrangement, we generated dotplots using Gepard (version 1.40) (Krumsiek et al. 2007) with the clade-A2 isolate E0139 as a reference.

### Characterization of fully assembled plasmids

Contigs derived from hybrid assembly were labeled either as chromosome or plasmid based on sequence length and circularization signatures. Contigs were categorized as plasmid if they presented circularization signatures and a sequence length smaller than 350 kbp. Putative plasmids smaller than 350 kbp and lacking circularization signatures were not considered for further analysis. Rapid annotation by Prokka (version 1.12) (Seemann 2014) allowed us to discard four putative circular phage sequences.

We used Abricate (version 0.8.2) to query (> 80% identity & > 60% coverage) our set of complete plasmid sequences (n = 305) versus a curated database of known replication initiator and relaxases proteins from *Enterococcus* (Jensen et al. 2010; Clewell et al. 2014). Replicon sequences with a lower identity (< 80% ID) were classified as Rip-like.

Complete plasmid sequences were clustered using Mash (k = 21, s = 1000 ; version 1.1) (Ondov et al. 2016) and the resulting distance matrix was clustered using the hclust function (method = ‘ward.D2’) provided in R package stats (version 3.4.4). Dendrogram visualization and metadata associated to plasmid isolates was displayed using ggtree (version 1.13.3) (Yu et al. 2017).

### Unraveling the plasmidome content of E. faecium using a machine-learning approach

To determine the plasmidome content of the remaining 1,582 isolates, we used mlplasmids which provides a support-vector machine classifier which was trained and tested using complete genome sequences from *E. faecium* (Arredondo-Alonso et al. 2018). In short, short-read contigs derived from our completed genomes were mapped against the finished chromosomes and plasmids to obtain a short-read contigs dataset labeled either as chromosome- or plasmid-derived. This dataset was used to train and test five popular classifiers using pentamer frequencies as model features. The best performing supervised classifier, a support vector machine (svm), predicted plasmids with an accuracy of 0.95 and an F1 score of 0.92 on the test set (Arredondo-Alonso et al. 2018).

Mlplasmids (version 1.0.0) was run specifying: ‘Enterococcus faecium’ model and a minimum contig length of 1,000 bp. For further analysis, we discarded predicted contigs with a posterior probability lower than 0.7 of belonging to the assigned class (chromosome or plasmid). Differences in the number of chromosome- and plasmid-derived base-pairs predicted by mlplasmids between hospitalized patient isolates and other isolation sources were assessed using Kruskal Wallis test (significance threshold = 0.05) available in ggpubr package (version 0.1.7) (Kassambara 2017).

We calculated pairwise Mash distances (k = 21, s = 1,000 ; version 1.1) between isolates (n = 1,640) uniquely considering plasmid-predicted contigs. We reconstructed plasmidome phylogeny with the bioNJ algorithm from Gascuel implemented in R ape package (version 5.1) using computed Mash distances (Gascuel 1997; Paradis et al. 2004). The resulting phylogenetic tree was midrooted using the midpoint function in the R phangorn package (version 2.4.0) (Schliep 2011). To improve the resolution of the bioNJ tree, we observed the distribution of the computed Mash distances and fitted a Gamma distribution using the fitdist function (distr = “gamma”, method = “mle”) available in R fitdistrplus package (Delignette-Muller et al. 2015). We discarded isolates with an average pairwise mash distance superior to 0.12 which was calculated using the qgamma function (p = 0.9, shape = 2.344073, rate = 35.870449, lower.tail = TRUE) in R stats package (version 3.4.4). All remaining isolates (n = 1,607) were used to reconstruct the plasmidome phylogeny.

We used the function NbClust (method = ‘ward.D2’, index = ‘silhouette’) available in R NbClust package (version 3.0) (Charrad et al. 2013) to evaluate an optimal number of clusters derived from pairwise Mash distances. We computed hierarchical clustering using the hcut function (method = ‘ward.D2’, isdiss = TRUE, k = 26) and cut the resulting dendrogram specifying 26 clusters. For each resulting cluster, we uniquely defined plasmid populations (n = 9) based on two criteria: i) clusters with more than 50 isolates and ii) an average silhouette width greater than 0.3.

Correlation of plasmid populations and isolation sources was determined using a one-sided Fisher-exact test (alternative = ‘greater’) from the fisher.test function (R stats package version 3.4.4) and naive p-values were adjusted using benjamini-hochberg (BH) method implemented in p.adjust function (R stats package, version 3.4.4). We considered an adjusted p-value threshold of 0.05 to determine enrichment of isolation sources for specific plasmid populations.

We incorporated metadata and plasmid population information into plasmid bioNJ and *E. faecium* core-genome tree using the R ggtree package (version 1.13.3).

To further investigate the influence of plasmids on *E. faecium* phylogeny, we again used Mash (k = 21, s = 1000) (Ondov et al. 2016) but on three different input data: i) plasmid-predicted contigs, ii) chromosome-predicted contigs and iii) whole-genome assembly. Mash distances were transformed into several bioNJ trees with the bionj function from ape package (version 5.1) (Paradis et al. 2004). Resulting trees were midpoint rooted using the midpoint function in the R phangorn package (version 2.4.0) and compared against our defined ‘benchmark’ *E. faecium* core genome tree. We performed pairwise comparisons of all trees using the Kendall and Colijn (KC) metric considering differences only between topology of the trees (*λ* = 0) and a balanced measure between branch and topology of the trees (*λ* = 0.5) (Jombart et al. 2017).

### Contribution of genomic components on host specificity

To evaluate the contribution of genomic components on host specificity, we considered three different inputs: i) Mash pairwise distances from whole-genome contigs, ii) Mash pairwise distances from chromosome-derived contigs and iii) Mash pairwise distances from plasmid-derived contigs. Pairwise distances were scaled using the scale function (scale = TRUE, center = TRUE) from R stats package (version 3.4.4). For each isolation source (hospitalized patient, dog, poultry, pig and non-hospitalized person), we used a bootstrap approach (100 iterations) to calculate the average pairwise distances of 50 randomly isolates belonging to the following combinations: i) pairs of isolates belonging to the same host (within-host group), ii) pairs of isolates belonging to different hosts (between-host group) and iii) pairs of isolates belonging to random isolation sources (random group). This random group consisted of an artificial group in which we merged 50 randomly isolates belonging to any of the five isolation sources after sampling 100 isolates from each of the sources to avoid overrepresentation of hospitalized patient isolates. This random group was used to statistically assess whether the distribution of pairwise distances belonging to within-host and between-host groups differed from random pairwise distances. We used a one-way anova test (aov function, R stats package version 3.4.4). and computed differences in the observed means using TukeyHSD function available in R stats package (version 3.4.4). A significant (adjusted *P* < 0.05) positive and negative observe difference mean was considered as an indication of host adaptation similarity and dissimilarity respectively.

### Scoary analysis to determine plasmid-derived orthologous gene enrichment

To observe plasmid OGs genes enriched in *E. faecium* hosts, plasmid-predicted contigs were annotated using Prokka (version 1.12) (Seemann 2014) specifying the *Enterococcus* database, and orthologous groups (pan-plasmidome) were estimated using Roary (version 3.8) (Page et al. 2015), splitting paralogues and defining a threshold of 95% amino-acid level similarity to cluster protein sequences. We ran Scoary (-p 1E-5, -c BH --no_pairwise, --collapse) (version 1.6.16) (Brynildsrud et al. 2016). We used recommended Scoary parameters (https://github.com/AdmiralenOla/Scoary) to observe plasmid genes overrepresented in specific host groups. Sensitivity and specificity (> 80%) reported values were used to highlight group of genes strongly associated to particular hosts.

### Co-evolution analysis

A core-genome alignment (1.1 MB) was generated using the Harvest suite v1.1.2, including the 1,644 Clade A isolates and the complete *E. faecium* AUS0004 (accession number CP003351) as reference (Lam et al. 2012). SuperDCA was performed with the default setting as described by Puranen et al. (Puranen et al. 2018) using the alignment of 1,644 isolates which was unfiltered for recombination events.

### Code availability

Complete code used to generate the analysis reported in our manuscript is publicly available under the following gitlab repository [https://gitlab.com/sirarredondo/efaecium_population].

### Data access

Illumina NextSeq500/MiSeq reads of the 1,644 E. faecium isolates used in this study have been deposited in the following European Nucleotide Archive (ENA) public project: PRJEB28495. Oxford Nanopore Technologies MinION reads used to complete the 62 E. faecium genomes are available under the following figshare projects: 10.6084/m9.figshare.7046804, 10.6084/m9.figshare.7047686. Hybrid assemblies generated by Unicycler (v.0.4.1) are available under the ENA project: PRJEB28495 and under the following gitlab repository https://gitlab.com/sirarredondo/efaecium_population/tree/master/Files/Unicycler_assemblies Raw core-genome alignment (1.1 MB, Harvest suite v1.1.2), including the 1,644 Clade A isolates and the complete E. faecium AUS0004 (accession number CP003351) as reference is available under the following gitlab repository https://gitlab.com/sirarredondo/efaecium_population/tree/master/Files/Harvest_suite Exploratory analysis of our data and metadata set is available under the following microreact project https://microreact.org/project/BJKGTJPTQ.

## Acknowledgments

SA and RJLW: This study was supported by the Joint Programming Initiative in Antimicrobial Resistance (JPIAMR Third call, STARCS, JPIAMR2016-AC16/00039).

J C was funded by the European Research Council (grant no. 742158).

WS was supported by a Royal Society Wolfson Research Merit Award.

PM: This work was funded by the Academy of Finland (grants no. 286607 and 294015).

Author contributions: JC and RW initiated and supervised the project. JB and MR performed the genomic DNA extraction, genome sequencing and assemblies. SA, JT, AS, AM, SP, JP, SK and MP performed analysis. SA, JT, AS, JC and RW wrote the manuscript with input from AM and WS

## Disclosure declaration

The authors declare no conflicts of interest.

