## Supplemental Material for "Genomes of a major nosocomial pathogen *Enterococcus faecium* are shaped by adaptive evolution of the chromosome and plasmidome"

### Table of Contents:

**Supplemental Results:**

- S1 Characterization of complete plasmid sequences obtained by ONT sequencing
- S2 The plasmidome of *E. faecium* differs substantially between hosts
- S3 Plasmid cluster of orthologous genes (OG) groups
- S4 Detailed description of co-evolving genomic regions with linked SNPs

**Supplemental Methods:**

- S1 Illumina sequencing
- S2 WGS short-read assemblies
- S3 Selection of isolates to sequence by ONT
- S4 ONT sequencing
- S5 Assembly of ONT sequenced isolates

**Supplemental Figures:**

- S1. RAxML tree based on 859 core genes in 40 clade B isolates. Nodes represent the source with indication of presence of vancomycin resistance gene, vanA (pink) or vanB (green).
- S2. Dotplots of chromosomal rearrangements observed in our set of 48 complete chromosome sequences using strain E0139 (Clade A2) as reference.
- S3 Core genome tree based on 1,644 strains with the three metadata panels: i) distribution of genomic rearrangements types among 38 complete

chromosomes, ii) indication of strains lacking a genomic rearrangement among 10 complete chromosomes and draft genomes and iii) indication of strains with insertion of a carbohydrate transport system encoding genomic island or a phosphotransferase system encoding genomic island.

- S4 Overview of the complete plasmid sequences (n = 294) with an associated replication initiator gene (RIP).
- S5 Maximizing resolution of bioNJ plasmid-based phylogeny.
- S6 Definition of plasmid populations. A) Plasmid dissimilarity matrix of pairwise Mash distances was clustered using hierarchical clustering (ward.D2).
- S7 Phylogenies comparison (chromosome-predicted, whole-genome and plasmid-predicted)
- S8 Distribution of orthologous genes (OG) groups based on isolation source.
- S9 Differences in the observed means of average pairwise distances when comparing within-host and between-host groups against our defined random group of isolates.
- S10 Circos plots of the core genome representing linked positions for ten loci with the highest number of linked SNPs.

### Supplemental Results

#### S1. Characterization of complete plasmid sequences obtained by ONT sequencing

The completely sequenced plasmids were first characterized based on their replication initiator proteins (RIP). Most of our complete plasmid sequences contained a RepA_N initiator sequence (n = 82) with similarity versus RIP sequences described in pLG1 megaplasmid (n = 55, accession number ADO66907) and the non-conjugative pRUM plasmid (n = 27, accession number NP_863172). RepA_N family was found in large plasmids (mean = 155.2 kbp) (Supplemental Fig. S4B), occasionally associated with other RIP sequences (n = 15, freq = 0.18) (Supplemental Fig. S4A) and present in hospitalized patients (n = 63), dog (n = 12), pig (n = 3), non-hospitalized persons (n = 2) and chicken isolates (n = 2). We also identified plasmids containing RepA_N-like (<80% identity) initiators (n = 20) in medium plasmids (mean 53.9 kbp).

Also the Inc18 family was ubiquitous in our collection of plasmid sequences (n = 57) and present in plasmids with a medium size (mean = 44.7 kbp) (Supplemental Figures S4B and S4C). Plasmids bearing Inc18 family showed higher levels of mosaicism than RepA_N plasmids and were frequently present in multireplicon plasmids (n = 30 ; freq = 0.53). In this study, we mainly found Inc18 sequences with similarity to the initiator sequence from pRE25 plasmid (n = 44, accession number Q9AL28) which was originally identified in *E. faecalis* from a raw-fermented sausage and associated with multiple antibiotic resistance genes (Teuber et al. 2003). We identified Inc18 plasmids in isolates from hospitalized patients (n = 49), dog (n = 7), chicken (n = 2), pig (n = 1) and non-hospitalized persons (n = 1).

The Rep_3 family was mostly found on small plasmids (n = 56, mean plasmid length = 10.8 kbp) (Supplemental Figures S4B and S4C) and rarely present in multireplicon plasmids (n = 6, freq = 0.11.) (Supplemental Fig. S4A). We found Rep_3 sequences with similarity to other small theta-replicating plasmids such as *Enterococcus durans* pGL (n = 22, accession number ADW93773) or *E. faecium* p200B (n = 8, accession number BAF44066). In contrast to RepA_N and Inc18 families, we uniquely found Rep_3 sequences in isolates from hospitalized patients (n = 49), dogs (n = 5) and chickens (n = 2). We detected a high number of plasmids containing Rep_3-like (<80% identity) initiator sequences (n = 54, mean pl. length = 17.1 kbp) and were occasionally associated to other rip families (n = 19, freq = 0.35).

Rep_trans family (n = 24) was mainly identified in plasmids with a medium size (mean = 16.36 kbp) (Supplemental Fig. S4B) and occasionally present in multireplicon plasmids (n = 9, freq = 0.38) (Supplemental Fig. S4A). We mainly found similarity with Rep_trans sequence from pRI1 (n = 16, accession number YP_001672021), a small cryptic mobilizable *E. faecium* plasmid from human and animal origin (Garcia-Migura et al. 2009). Plasmids bearing Rep_trans family were present in hospitalized patients (n = 19), dog (n = 3) and chicken (n = 2) isolates. We also characterised Rep_trans-like (<80% identity) sequences (n = 36, mean pl. length = 20.8 kbp) with a similar frequency of being associated to other rip sequences (n = 11, freq = 0.31).

Rep_2-like (<80% identity) sequences (n = 10) were present in small-medium plasmids (24.0 kbp), frequently present in multireplicon plasmids (n = 6, freq = 0.6), similar to pJB01 (n = 9, accession number YP_138502) (Kim et al. 2006) and only present in hospitalized (n = 8) and dog isolates (n = 2).

Finally, we described another known *Enterococcus* Rip family corresponding to Rep_1 (n = 8, mean = 33.4 kbp). This Rip group was uniquely present in multireplicon plasmids in association with RepA_N and Inc18 families (n = 8, freq = 1.0). All Rep_1 sequences had similarity to a previously described non-functional Rep of the non-conjugative pAMα1 (accession number NP_863351) plasmid from *E.*

*faecalis* (Francia and Clewell 2002). Our findings were in accordance with previous reports suggesting Rep_1 family may not be functional and other rip initiators are required for plasmid replication (Clewell et al. 2014). Plasmids carrying Rep_1 initiators were found in patients (n = 5), dogs (n = 2) and non-hospitalized person isolates (n = 1). We additionally found Rep_1-like (<80% identity) sequences (n = 6) present in small plasmids (mean = 4.0 kbp), not associated with other replication initiator families and with similarity to Rep_1 sequences from *E. faecalis* pTEF1 (n = 6, accession number NP_816941) and *E. faecium* pNJAKD (n = 2, accession number YP_004747351).

MOB_P family was the most predominant relaxase family (n = 124) (Supplemental Fig. S4C) and present in plasmids with a single RepA_N (n = 44) or Rep_3 (n = 46) initiator sequence (Supplemental Fig. S4A). This relaxase family was also found in RepA_N-like (n = 15), Inc18 (n = 8), and other multireplicon plasmids (n = 11). MOB_V family was mainly found in plasmids carrying a single Rep_trans-like (n = 11), Rep_1-like (n = 5), Rep_trans (n = 2) families and multireplicon plasmids (n = 11) containing different combinations of Inc18, Rep_1, RepA_N and Rep_1 families (Supplemental Fig. S4A). MOB_T family was identified in multireplicon plasmids containing a Rep_trans-like group and several combinations of Inc18, RepA_N an Rep_1 sequences (n = 6). MOB_C was uniquely found in RepA_N plasmids (n = 2) including a multireplicon plasmid (RepA_N, Rep_trans and Rep_trans-like) (Supplemental Fig. S4A).

#### S2. The plasmidome of *E. faecium* differs substantially between hosts

Plasmid subpopulations (n = 9) were visualized in both the plasmid-bioNJ (Fig. 5A) and ML core-genome tree (Fig. 5B). We confirmed that plasmid subpopulations 1 (non-hospital and pig-related), population 10 (hospital-related), population 14 (dog-related), population 15 (hospital-related) and population 23 (hospital-related) were driven by horizontal transmission as these populations were adjacent to each other in the plasmid-bioNJ tree (Fig. 5A) but were distant in the ML core-genome tree (Fig. 5B) . Furthermore, we observed that plasmid population 5 (chicken-related), population 19 (hospital-related), population 25 (hospital-related) were mostly driven by vertical transmission as we observed co-clustering of the isolates from these populations in both ML core-genome and plasmid-NJ tree (Fig. 5).

**S3. Plasmid cluster of orthologous genes (OG) groups analysis**

Following the observation that the plasmidome of *E. faecium* isolates differed substantially between hosts, we next used Scoary (Brynildsrud et al. 2016) to determine cluster of orthologous genes (COG) groups and predicted which OGs plasmid sequences were enriched for a particular *E. faecium* host. For each host, we highlighted OGs with a specificity and sensitivity higher than 80%, which indicated that these genes were overrepresented for a particular host and underrepresented for the rest.

In isolates from hospitalized patients, 589 OGs had an Odds-ratio (OR) higher than 1 indicating a large pool of plasmid genes that were enriched in clinical *E. faecium* isolates (Supplemental Fig. S8). Of these OGs only 15 had a specificity of > 80% for hospitalized patients and a sensitivity > 80% . These 15 OGs include antimicrobial resistance genes (e.g. *aacA-aphD*), genes encoding efflux ABC transporters (permease proteins), DNA ligase (*ligA)*, a two-component system formed by a sensor histidine kinase (HK) and a DNA-binding response regulator (RR) and a serine recombinase and seven genes encoding hypothetical proteins (Supplemental Table S6).

From these hypothetical proteins, we observed that OG ‘group_118’ encoded for a protein of 783 aa with two domains: a peptidoglycan binding domain (pgdb) (coordinates 69-173) and a GH25_BacA-like domain (coordinates 277-485). The first domain is part of the PGBD-like superfamily (IPR036365) that have a general peptidoglycan binding function. The second domain belongs to the glycoside hydrolase superfamily (IPR017853) which can hydrolyse the glycosidic bond between carbohydrates.

We searched our protein sequence against BacA homologues, a Bac41-like gene which have been described as plasmid-encoded bacteriocin in *E. faecalis (Kurushima et al. 2016),* BacA homologues are splitted into five different clades. In our study, we observed a perfect match (blastp, e-value = 0.0, identity = 99%) between OG 'group_118' and EOK45589 which belongs to clade IV variant. We could not identify other Bac41-like genes in the adjacent areas of BacA, which is in accordance with the findings of BacA clade IV described by Kurushima et al 2016. Furthermore the authors argue about the functionality of this BacA homologue since they showed that the presence of BacL1 (another Bac41-like gene) is required for bacteriolysin activity. However, the high prevalence of this BacA homologue in *E. faecium* clinical isolates may suggest that the mechanism of this bacteriolysin have diverged for *E. faecium*. Kurushima et al. 2016 described that BacA can act as a more evolved toxin-antitoxin system in which not only daughter cells but also cells from the same generation not bearing BacA plasmid are excluded. Furthermore, authors showed that plasmid dissemination was more prominent under conditions of *E. faecium* populations fluctuations since Bac41 activity exclusively affects dividing cells.

We observed 155 plasmid genes enriched in poultry isolates plasmid genes (OR > 1) (Supplemental Fig. S8). Of these, 16 plasmid genes had a sensitivity and specificity > 80%), which included metabolism genes involved in glucose transport (*gdh)* or small solutes transport (*yhjX)* , OGs encoding for antibiotic resistance proteins like a TetR family transcriptional regulator (OG group_431) or a choloylglycine hydrolase (OG group_1067), a putative tetronasin resistance protein (OG group_699) or stress response proteins (*cspC*) involved in cold shock response (Supplemental Table S6).

We found that 100 plasmid genes were enriched in pig isolates (OR > 1) (Supplemental Fig. S8), of which 16 had a specificity and sensitivity > 80%. These OGs included several genes involved in copper resistance which was a growth-promoting agent commonly used in piglets (Hasman et al. 2006). We manually annotated copper genes designated by Prokka using the previously described *E. faecium tcrYAZB* operon (Supplemental Table S6).

Finally, 87 and 64 enriched genes (OR > 1) in dog and non-hospitalized person isolates, respectively but none of these gene had a specificity and sensitivity > 80%. This is most likely due to the fact that for the dogs a fraction of their plasmid genes is also found in other isolation sources and more likely with hospitalized patients while for the isolates from non-hospitalized persons this probably reflects the heterogeneity of this group.

**S4. Detailed description of co-evolving genomic regions with linked SNPs**

Because of the large number of significant couplings between SNP loci identified by SuperDCA, we generated a list of 10 loci with the highest number of linked SNPs and converted these loci into the genes and/or intergenic regions annotated in AUS0004 (Supplemental Fig. S10).

Loci with the highest number of linked SNPs involved two chromosomal regions, region-1 (*EFAU004_02173* to *EFAU004_02178*) and region-2 (*EFAU004_00665* to *EFAU004_00675*) (Fig. 7A). For the prediction of the biological functions of the proteins encoded by region-1 and region-2 we used BLAST. Locus *EFAU004_02173* in region-1 is predicted to encode a membrane protein with domains of unknown function (DUF), while *EFAU004_02175* was predicted to encode for a permease with unknown specificity. BLAST analysis revealed more specific putative biological functions for *EFAU004_02176*, putatively encoding a thioredoxin (Trx), *EFAU004_02177*, a transcriptional regulator Rex1 and *EFAU004_02178* a putative ABC transporter (Fig. 7B). The exact function of the ABC transporter (EFAU004_02178) is difficult to predict despite the presence of several domains. The protein contains two so-called AAA domains, and a leucine-zipper (bZIP) domain. Proteins with AAA domains are members of a conserved family of ATP-hydrolyzing proteins with all kind of activities in many cellular pathways, including replication, DNA and protein transport, transcriptional regulation, ribosome biogenesis, membrane fusion, and protein disaggregation or degradation (Elsholz et al. 2017). bZIP domains are known to be involved in transcriptional regulation, regulating transcription in stress conditions, like heat in Salmonella (Hurme et al. 1996, 1997), or abiotic stress in eukaryotes, e.g. plants (Alves et al. 2014). Furthermore, BLAST to non-Enterococcaceae revealed co-localization of a similar ABC transporter and *rexA* gene in other species like *Streptococcus pneumoniae*, *Streptococcus agalactiae*, *Listeria monocytogenes* and *Chlamydia trachomatis*, suggesting that this ABC transporter might also be involved in some kind of stress response.

Loci *EFAU004_00665* and *EFAU004_00670* in region-2 are predicted to have very similar domains and to encode for a NAD(P)H:quinone oxidoreductase. EFAU004_00670 contains transmembrane helices and is therefore likely membrane bound while EFAU004_00665 lacks these transmembrane helices and is therefore likely to be soluble. BLAST revealed that the EFAU004_00670 protein is very similar to YhdH of *Escherichia coli* (48% amino acid) and EFAU004_00665 to $Qor_{Ec}$ of *E. coli* (32% amino acid) and $Qor_{Tt}$ *Thermus thermophiles* (30% amino acid). In *Staphylococcus aureus*, expression of a gene cluster containing two *qor*-like genes was induced under oxidative stress conditions (Maruyama et al. 2003) . These two Qor proteins (SA1988 and SA1989) are predicted as soluble and SA1988 had amino acid 26% similarity with EFAU004_00665. Locus *EFAU004_00671* is predicted to encodes for a putative transcriptional regulator as it contains a helix-turn-helix motif, which may regulate the expression of *EFAU004_00672* to *EFAU004_00675*. These genes display similarity with a polyamine transport system as described for *Streptococcus pneumoniae* including an ATP binding protein PotA, 2 permeases PotB and PotC and a substrate binding protein PotD (Shah and Swiatlo 2008). Polyamines are polycationic molecules and are required for optimal growth in both eukaryotic and prokaryotic cells and are implicated in pathogenicity of *S. pneumoniae*.

### Supplementary Methods

#### S1. Illumina sequencing

Bacterial isolates were grown overnight (O/N) at 37°C on blood agar plates. Single colonies were picked up and grown O/N at 37°C with Brain Heart Infusion (BHI). Bacterial cell pellets were pretreated and incubated 1-4 hours with 180 µL of enzymatic lysis buffer. Subsequently, 0.75 mg proteinase K were added and incubated at 56°C until lysis completion. 20 µL of RNAse A (10mg/mL) were added and incubated for 5' at room-temperature (RT). Total DNA purification was performed using and following the protocol from NucleoSpin 96 Tissue Core Kit (Machery-Nagel), vacuum processing. DNA concentration was measured using Quant-it Picogreen (Thermo Fisher Scientific). Library preparation was carried out following Nextera DNA Library Prep Reference Guide. Finally, Nextera libraries were sequenced using Illumina NextSeq at USEQ, Utrecht, The Netherlands (http://www.useq.nl).

#### S2. WGS short-read assemblies

Illumina reads were trimmed using nesoni clip, part of the nesoni toolkit (version 0.132), with the following settings: '--adaptor-clip yes --match 10 --max-errors 1 --clip-ambiguous yes --quality 10 --length 90 --trim-start 0 --trim-end 0 --gzip no --out-separate yes pairs:'. Trimmed reads were then assembled into scaffolds using SPAdes (version 3.5.0) with default settings. Scaffolds with an average coverage lower than 10 and/or a length smaller than 500bp were removed from the assemblies.

#### S3. Selection of isolates to sequence by ONT

A fraction (n=62) of the total number of isolates was selected for long-read sequencing using Nanopore technology. We initially predicted the plasmid content of the isolates *in silico* using *PlasmidSPAdes* (version 3.8.2) which performs *de novo* assembly filtering out contigs with a coverage similar to the host chromosome coverage (Antipov et al. 2016). *Prokka* (version 1.12) was used to annotate the putative remaining plasmid contigs specifying the custom *Enterococcus* database provided (Seemann 2014). Orthologous clustered genes were estimated using *Roary* (version 3.8), splitting paralogues and defining a threshold of 95% amino-acid level similarity to cluster protein sequences (Page et al. 2015). This multi-dimensionality matrix was then reduced and visualized to two dimensions using the t-Distributed Stochastic Neighbor Embedding (*t-SNE*) (theta = 0.5, iterations = 1000, dims = 2) using the implementation provided in the *R* package *Rtsne* (version 0.13) (Maaten and Hinton 2008; Krijthe 2015). To avoid manual selection of the isolates, k-means function (iter.max = 1000) provided in the *R* package stats (version 3.4.4) was used to and allocated 50 centroids into the dimensionality reduced distribution given by *tSNE. Euclidean* distance of each isolate was calculated to extract the 50 isolates closest to each centroid.

To cover all plasmid replication genes not present in the first selection, 12 additional isolates were selected for Nanopore sequencing. This second selection was based on a reciprocal blast (blastx and tblastn, -evalue 1e-10) of the predicted plasmid orthologous genes against 76 previously described plasmid replication amino-acid sequences from the genus *Enterococcus* (Clewell et al. 2014). Isolates bearing plasmid replication genes not present in the first selection were sorted and selected based on highest number of orthologous genes.

#### S4. ONT sequencing

*E. faecium* selected isolates (n = 62) were grown O/N at 37°C on blood agar plates, then single colonies were picked up and grown with BHI at 37°C. Genomic DNA was extracted using the Wizard Genomic DNA purification kit (Promega) following manufacturer's instructions. Isolated DNA was sheared (4000 rpm, 2x120 seconds) using G-tubes (Covaris). Library preparation was performed using Ligation Sequencing Kit 1D (SQK-LSK108) with the Native Barcoding Kit 1D (EXP-NBD103). Genomic libraries were loaded onto R9.4 (FLO-MIN106) flowcells using the MinION device (Mk2). Libraries were basecalled using Metrichor workflows (Run 1 ,2, 3), Albacore 1.01 (Run 4, 5) and Albacore 1.1.0 (Run 6). ONT Sequencing and basecalling were conducted at USEQ, Utrecht, The Netherlands (http://www.useq.nl)

**S5. Assembly of ONT sequenced isolates**

Fastq files were obtained from base-called data using Poretools (version 0.6.0) except for Run6 in which fastq files were retrieved using Albacore (version 1.1.0). Distribution of read length and total number of reads were calculated using Bioawk (version 20110810, https://github.com/lh3/bioawk). We used Porechop (version 0.2.1, https://github.com/rrwick/Porechop) to trim reads and filter out chimeras from different bins specifying the flag "--discard_middle". Illumina reads were trimmed using seqtk (version 1.2-r94, https://github.com/lh3/seqtk) with the command "--trimfq" prior to assembly.

Hybrid assembly was performed using Unicycler (version 0.4.1), specifying "bold" mode (Wick et al. 2017). Briefly, Unicycler uses SPAdes (version 3.6.2) to create different assembly graphs based on different k-mer size only considering Illumina reads (Bankevich et al. 2012). The best assembly graph was selected by Unicycler based on number of dead-ends and contiguity. Next, all ONT reads were used to scaffold and solve the assembly graph. Additionally, we specified the same file as described above (section '*Selection of isolates to sequence by ONT*') containing 76 known plasmid replication sequences to rotate and change the 0-coordinate of circular replicons resulting from hybrid assembly (Clewell et al. 2014). Finally, Unicycler conducted several rounds of Pilon (version 1.22) to polish genome sequences using Illumina reads (Walker et al. 2014).

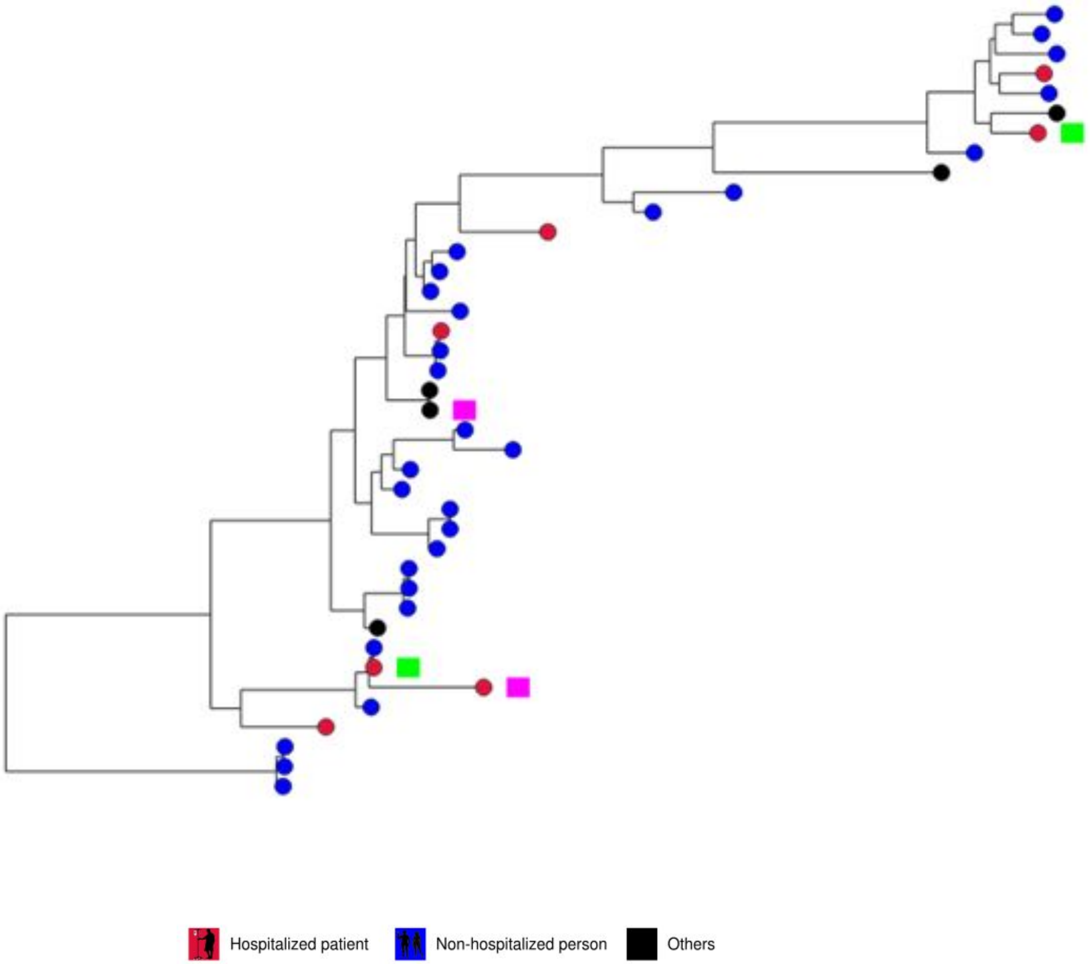


Supplemental Fig. S1. RAxML tree based on 859 core genes in 40 clade B isolates. Nodes represent the source with indication of presence of vancomycin resistance gene, vanA (pink) or vanB (green).

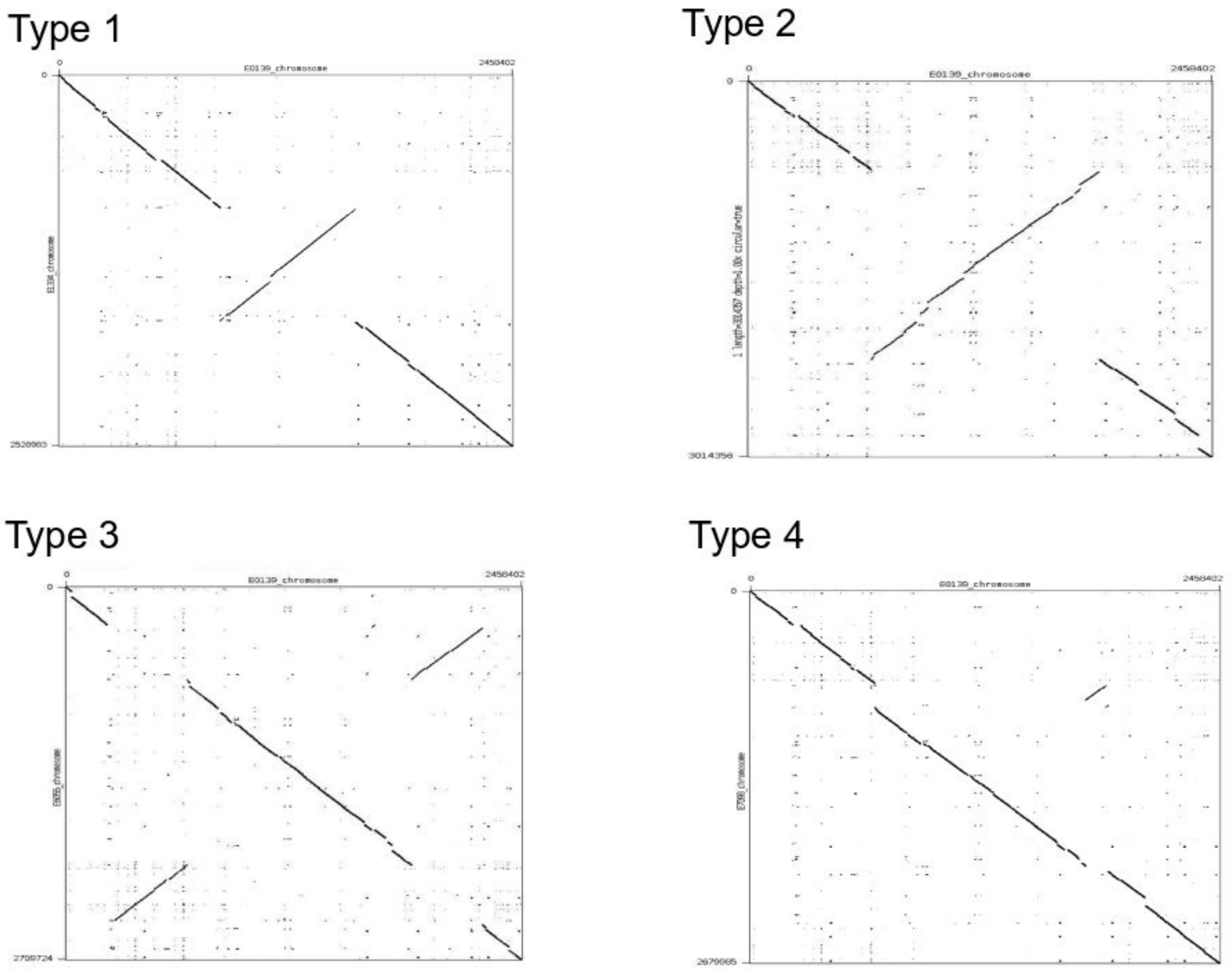


Supplemental Fig. S2. Dotplots of chromosomal rearrangements observed in our set of 48 complete chromosome sequences using strain E0139 (Clade A2) as reference. A, Chromosomal rearrangement Type 1 (E1334). B, Chromosomal rearrangement Type 2 (E8202). C, Chromosomal rearrangement type 3 (E6055). D, Chromosomal rearrangement type 4 (E7098).

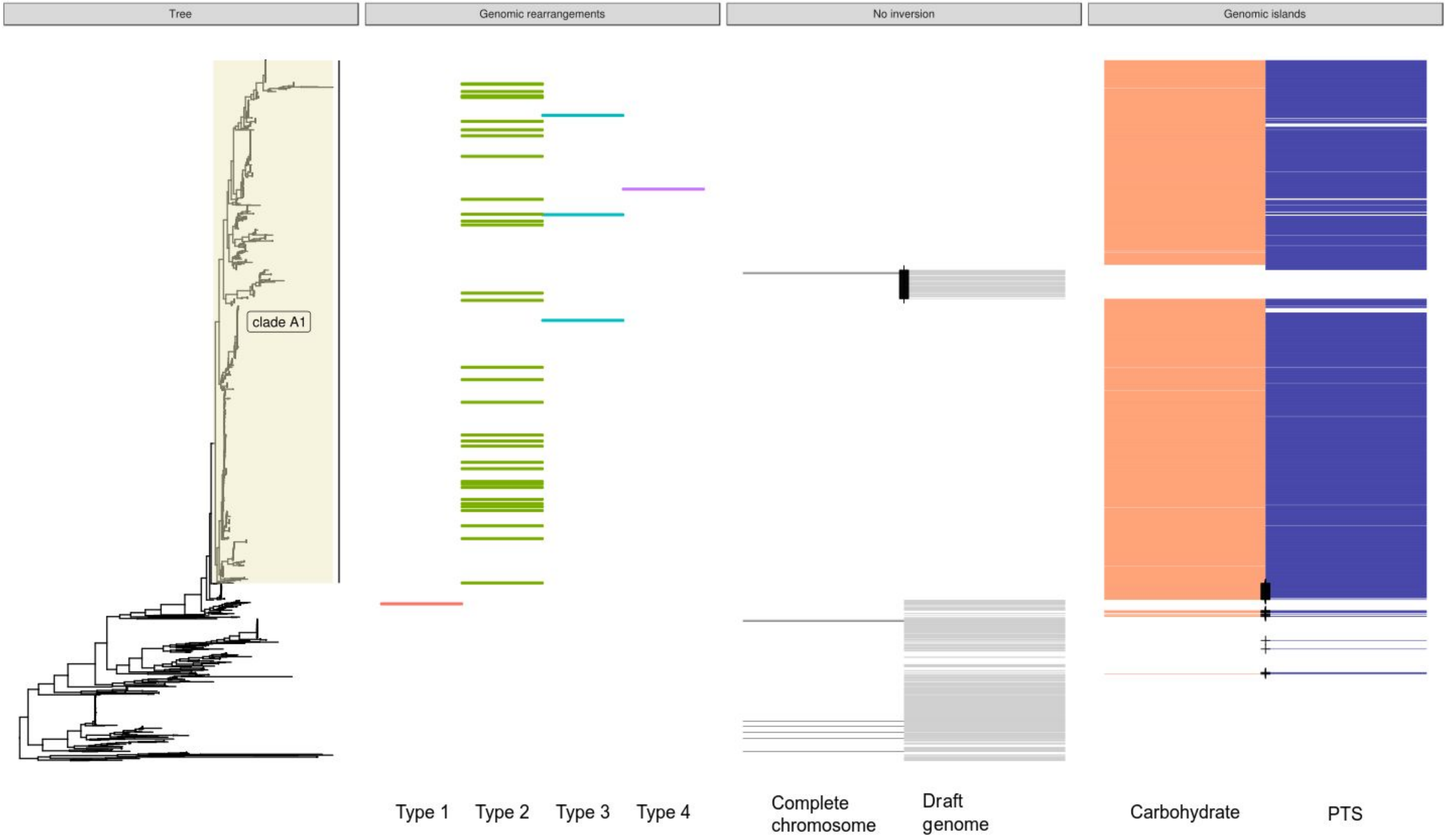


Supplemental Fig S3. Core genome tree based on 1,644 strains with the three metadata panels: i) distribution of genomic rearrangements types among 38 complete chromosomes, ii) indication of strains lacking a genomic rearrangement among 10 complete chromosomes and draft genomes and iii) indication of strains with insertion of a carbohydrate transport system encoding genomic island (orange)[33] or a phosphotransferase system encoding genomic island (purple)[34]. Black cross: indication of clade A1 dog isolates that lack the insertions/inversion or non-clade A1 dog isolates that do contain insertions/inversion.

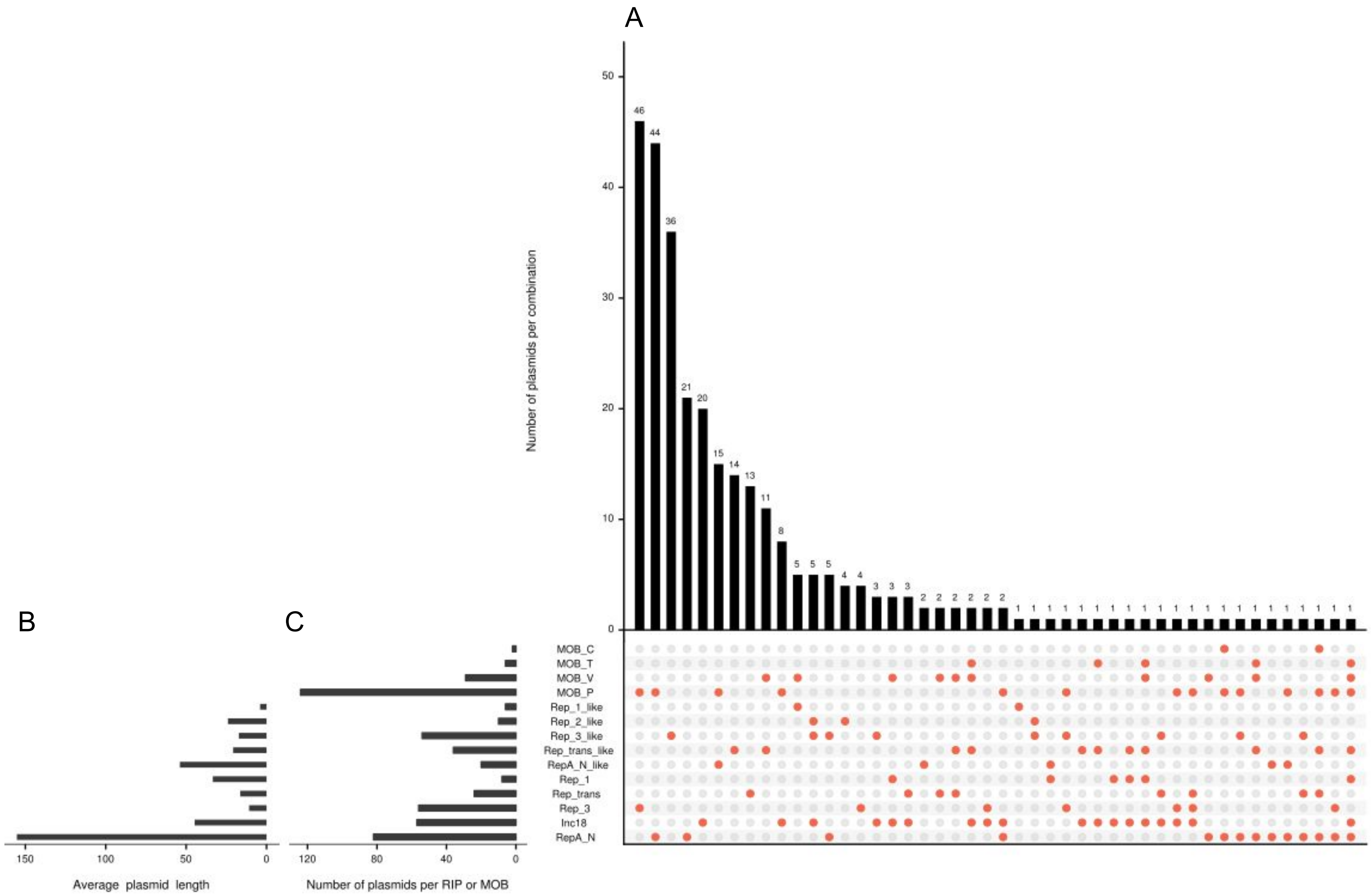


Supplemental Fig. S4. Overview of the complete plasmid sequences (n = 294) with an associated replication initiator gene (RIP). A) Histogram of the number of plasmids based on the combinations of RIP and MOB groups. B) Mean plasmid length (kbp) per RIP group. C) Number of plasmids per RIP or relaxase (MOB) group.

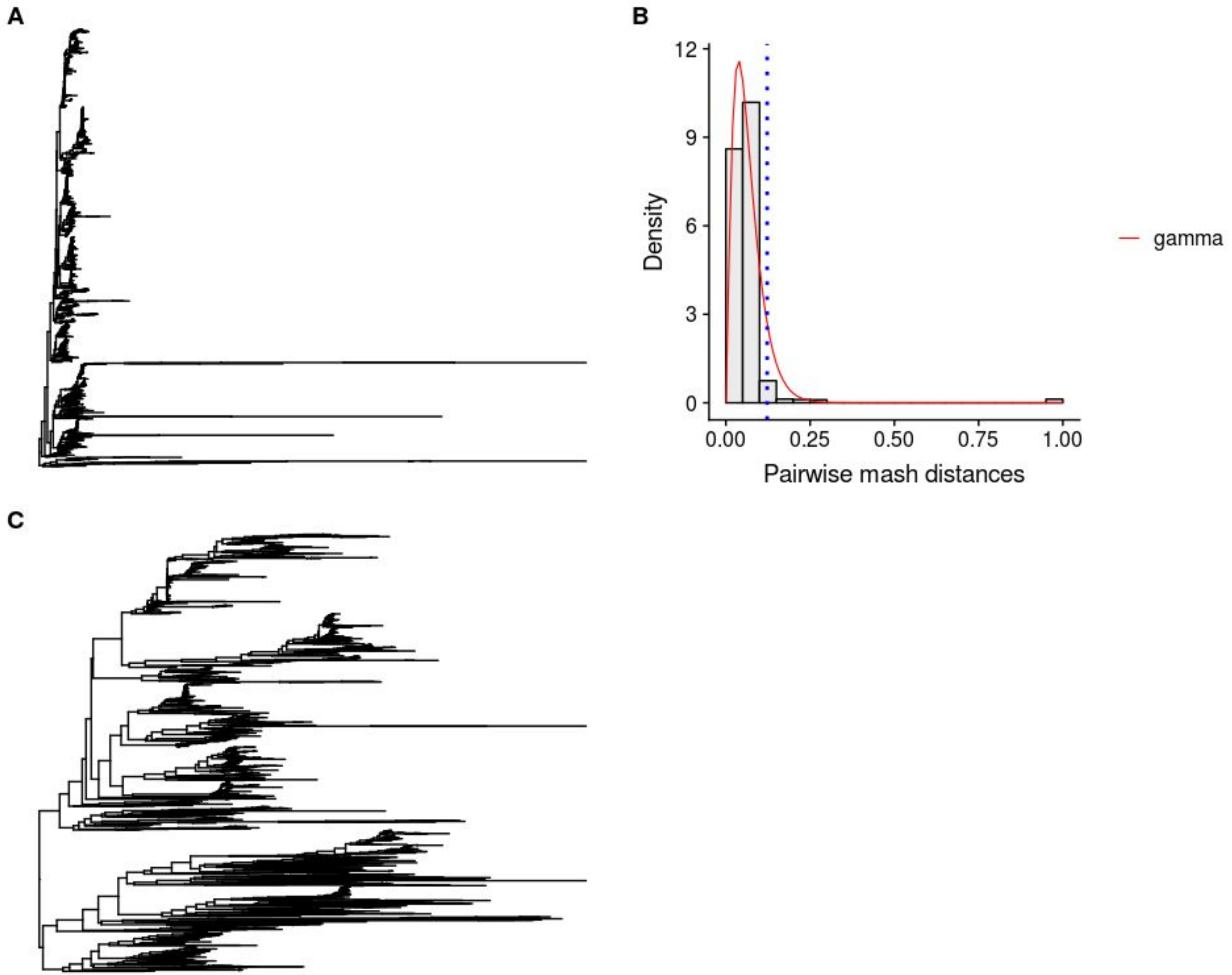


Supplemental Fig. S5. Maximizing resolution of bioNJ plasmid-based phylogeny. A) bioNJ tree of 1,639 isolates considering plasmid-predicted contigs by mlplasmids. B). Histogram against fitted density functions of pairwise Mash distances obtained by denscomp function (fitdistrplus R package). Vertical dash line indicates the Mash distance (0.1224967) used to filter out isolates (n = 37) with a higher average pairwise Mash distance. C) bioNJ plasmid-based phylogeny of 1,607 isolates after exclusion of 32 isolates.

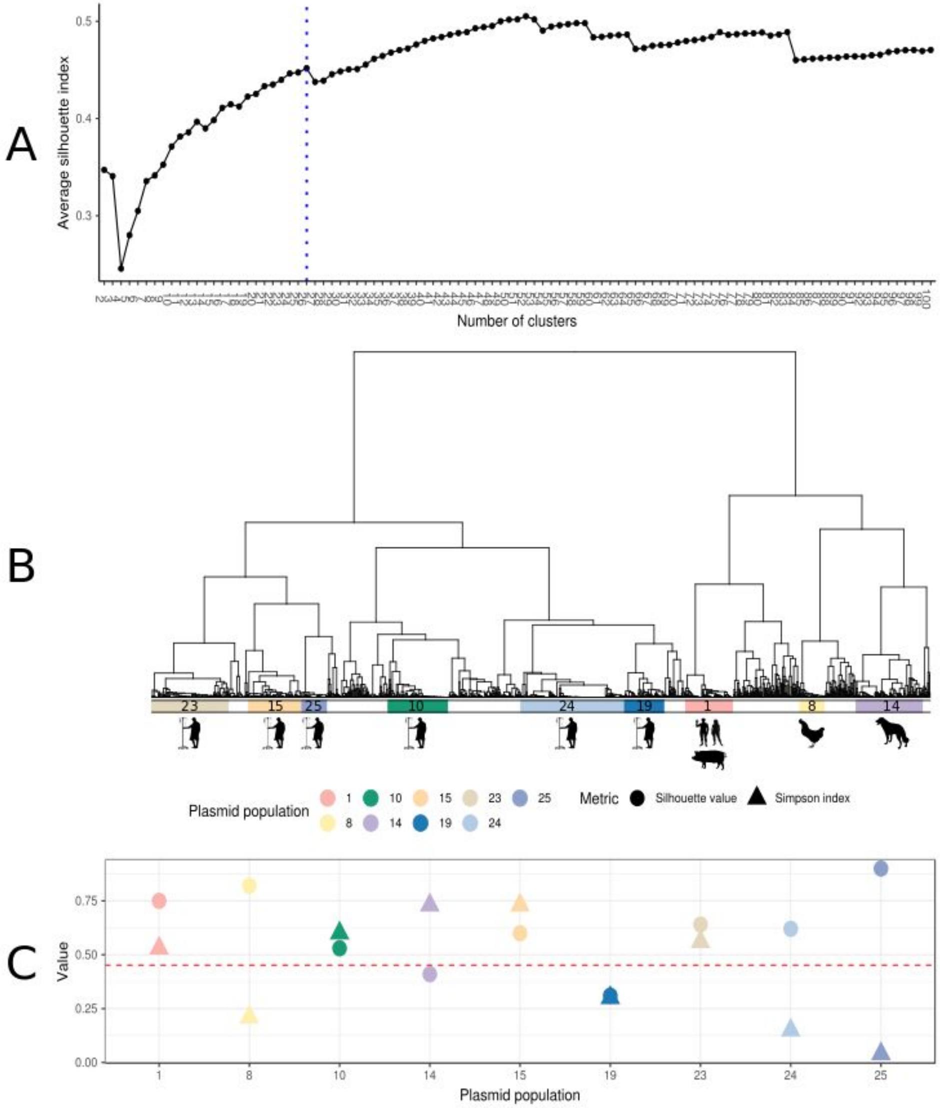


Supplemental Fig. S6. Definition of plasmid populations. A) Plasmid dissimilarity matrix of pairwise Mash distances was clustered using hierarchical clustering (ward.D2). Average silhouette index was computed based on the number of clusters (2–100). Vertical dash line corresponds to the clustering solution (k = 26) selected to cut the resulting dendrogram into 26 clusters. B) From these 26 clusters, we only defined plasmid populations (n = 9) if a particular cluster had a size larger than 50 isolates and an average silhouette index higher than 0.3. Each plasmid population was overrepresented at least with one isolation source. C) Simpson index (based on diversity of SC groups) and average silhouette index of each plasmid population. Horizontal dashed line indicates the average silhouette index of the selected clustering solution (k = 26, avg. silhouette index = 0.42).

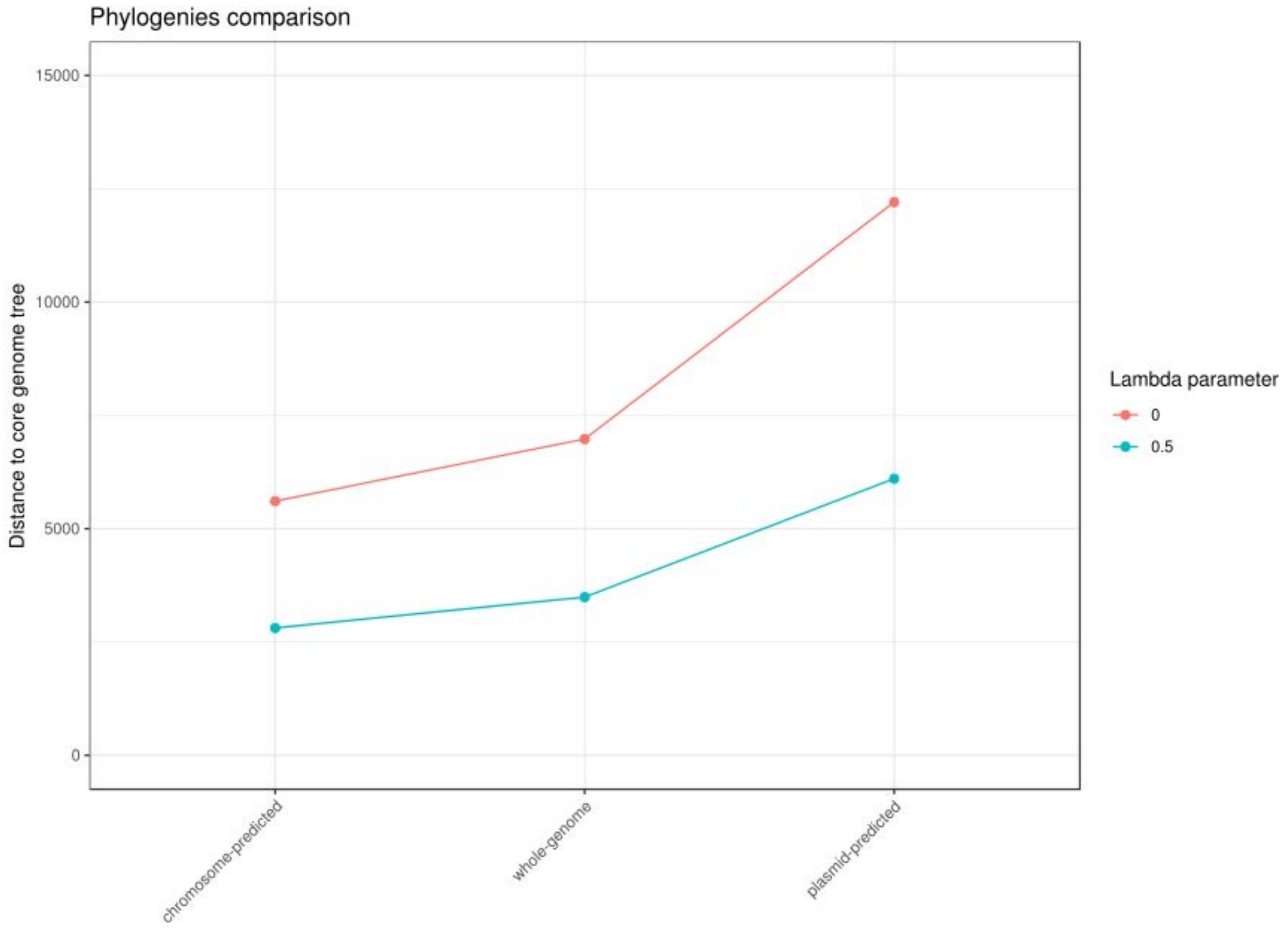


Supplemental Fig. S7. Phylogenies comparison (chromosome-predicted, whole-genome and plasmid-predicted) based on Kendall-Colijn (KC) metric considering differences only between topology of the trees ($\lambda$ = 0) and a balanced measure between branch and topology of the trees ($\lambda$ = 0.5) using as ground truth the core genome alignment.

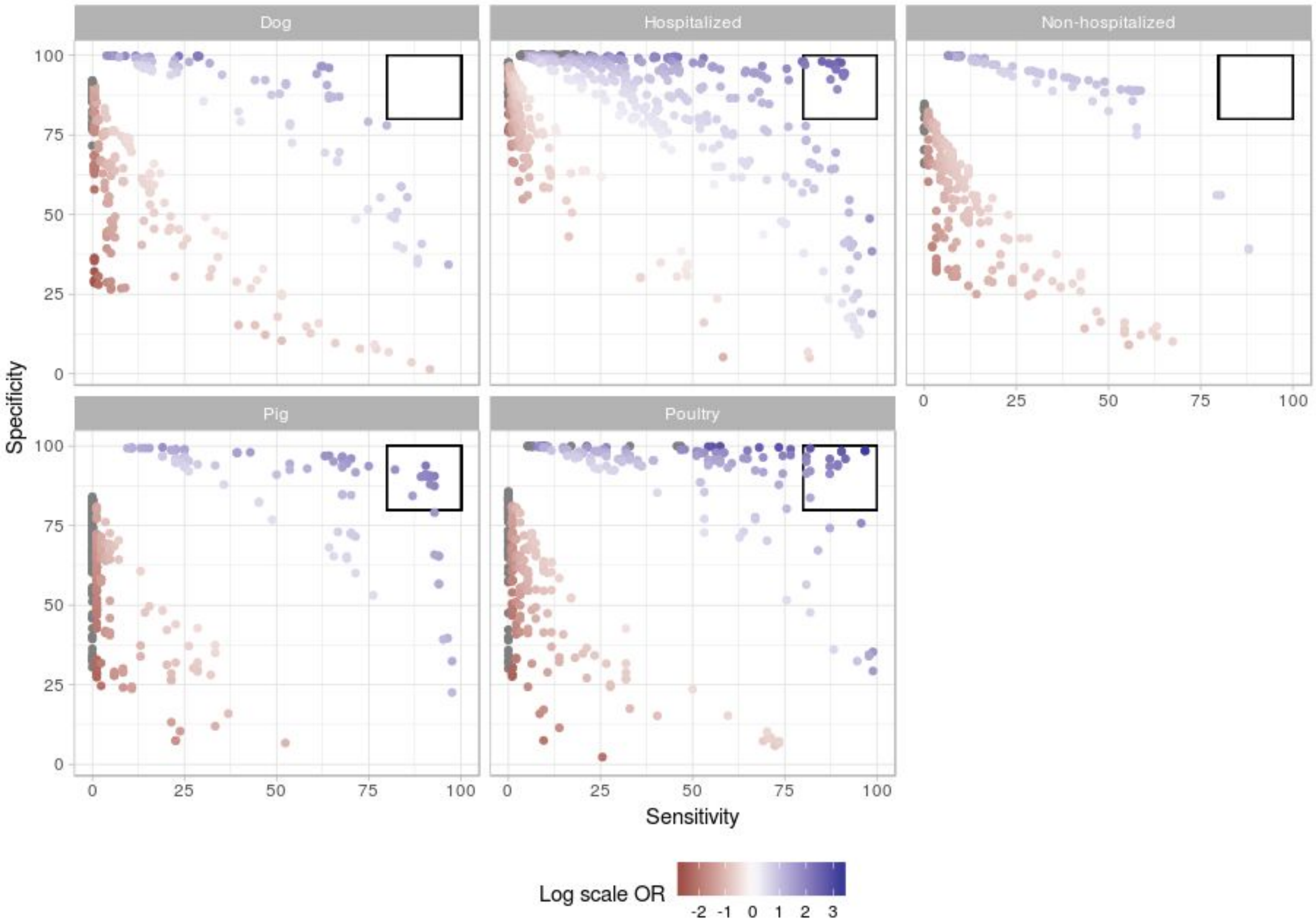


Supplemental Fig. S8. Distribution of orthologous genes (OG) groups based on isolation source. For each isolation source, Odds-ratio (OR) was transformed into a log scale (-2 to 3) to colour OG groups underrepresented (red) and overrepresented (blue). We further characterized OG groups (black square) with a specificity and sensitivity higher than 80%.

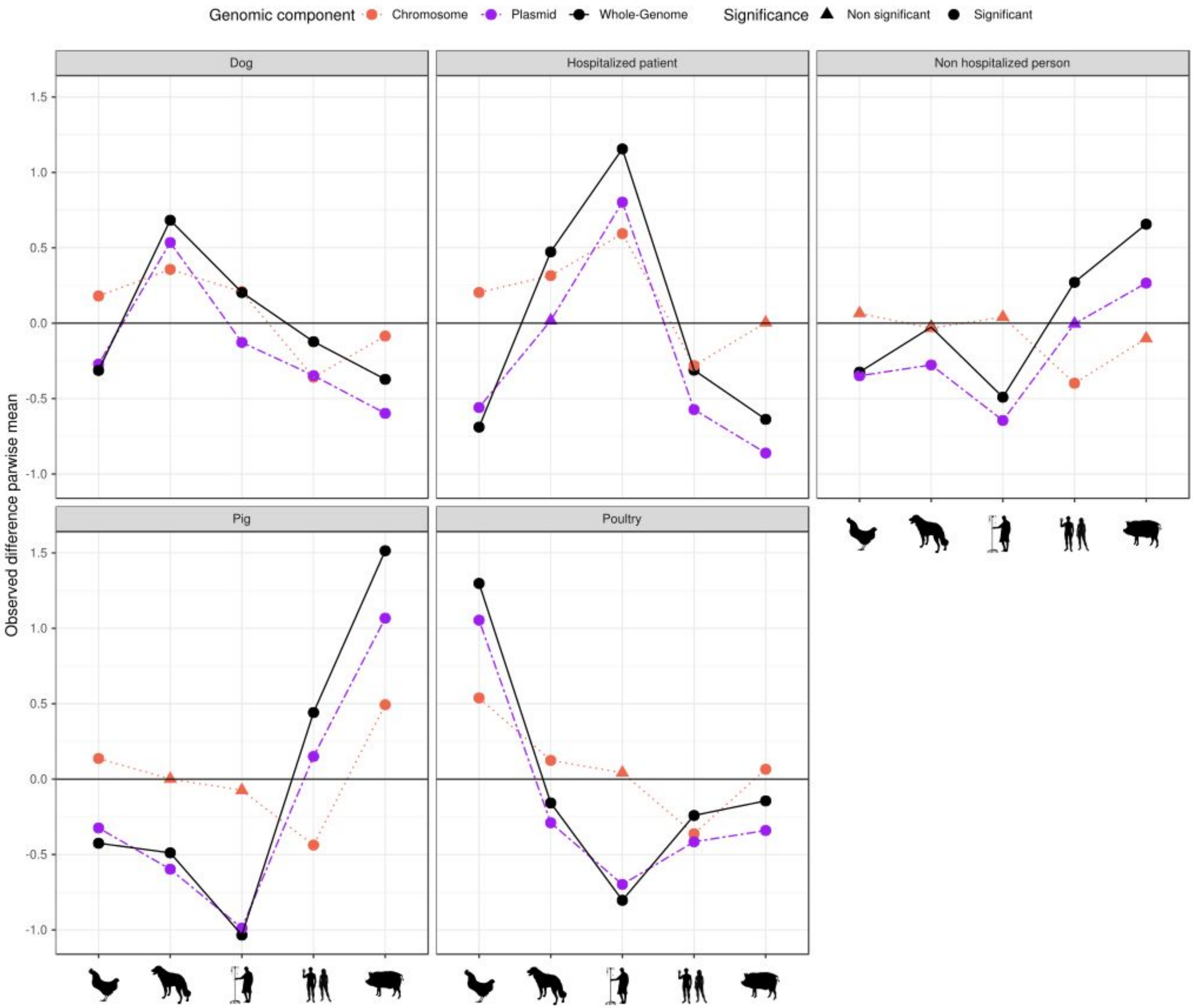


Supplemental Fig. S9. Differences in the observed means of average pairwise distances when comparing within-host and between-host groups against our defined random group of isolates. Each line corresponds to a different genomic component (whole genome - black solid, chromosome - dotted red, plasmid - dashed purple) and test significance is indicated based on shape (triangle - non significant, circle - significant).

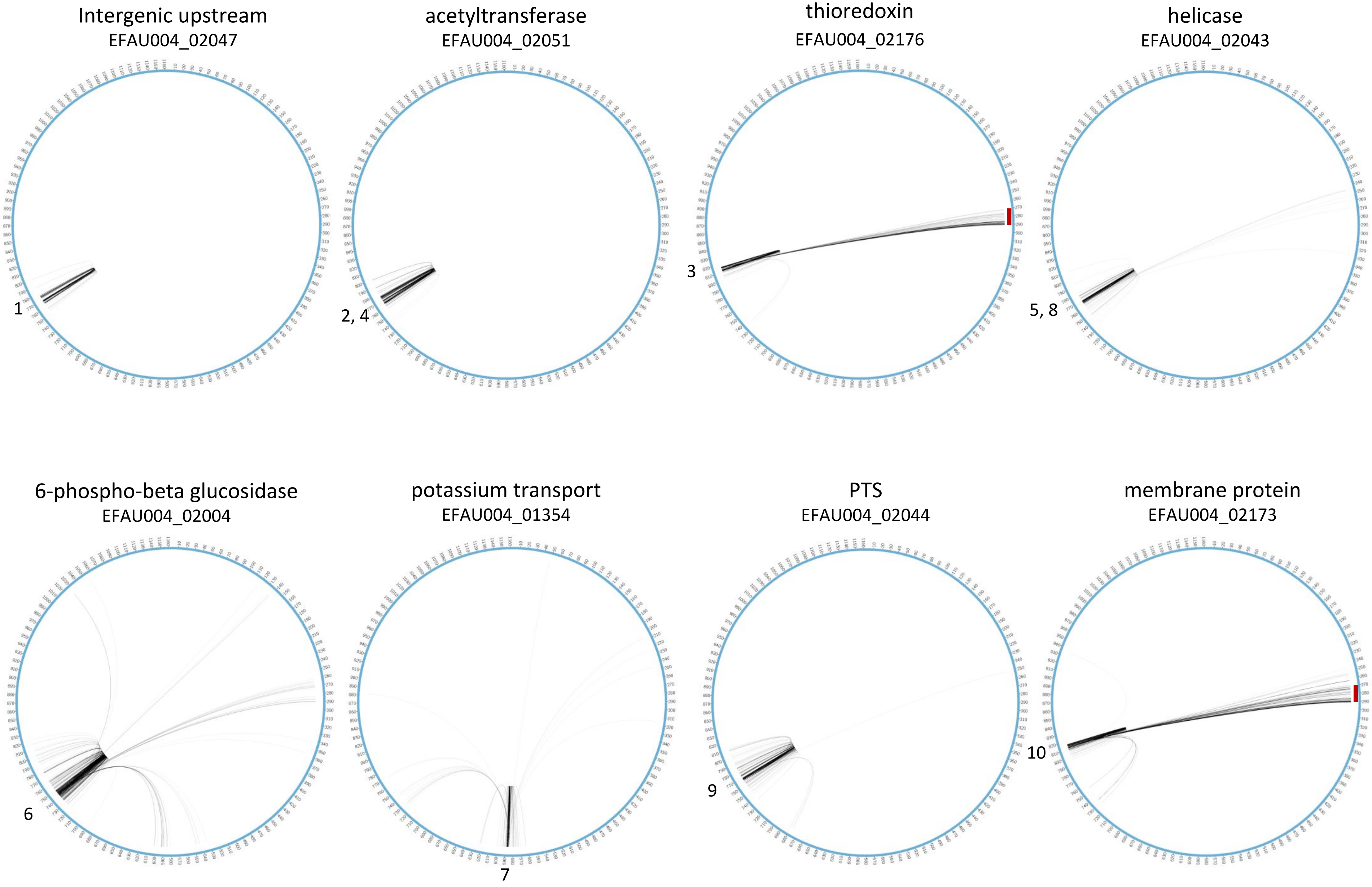


Supplemental Fig. S10. Circos plots of the core genome representing linked positions for ten loci with the highest number of linked SNPs. At the top of each plot, the converted position of loci in the AUS0004 genome.

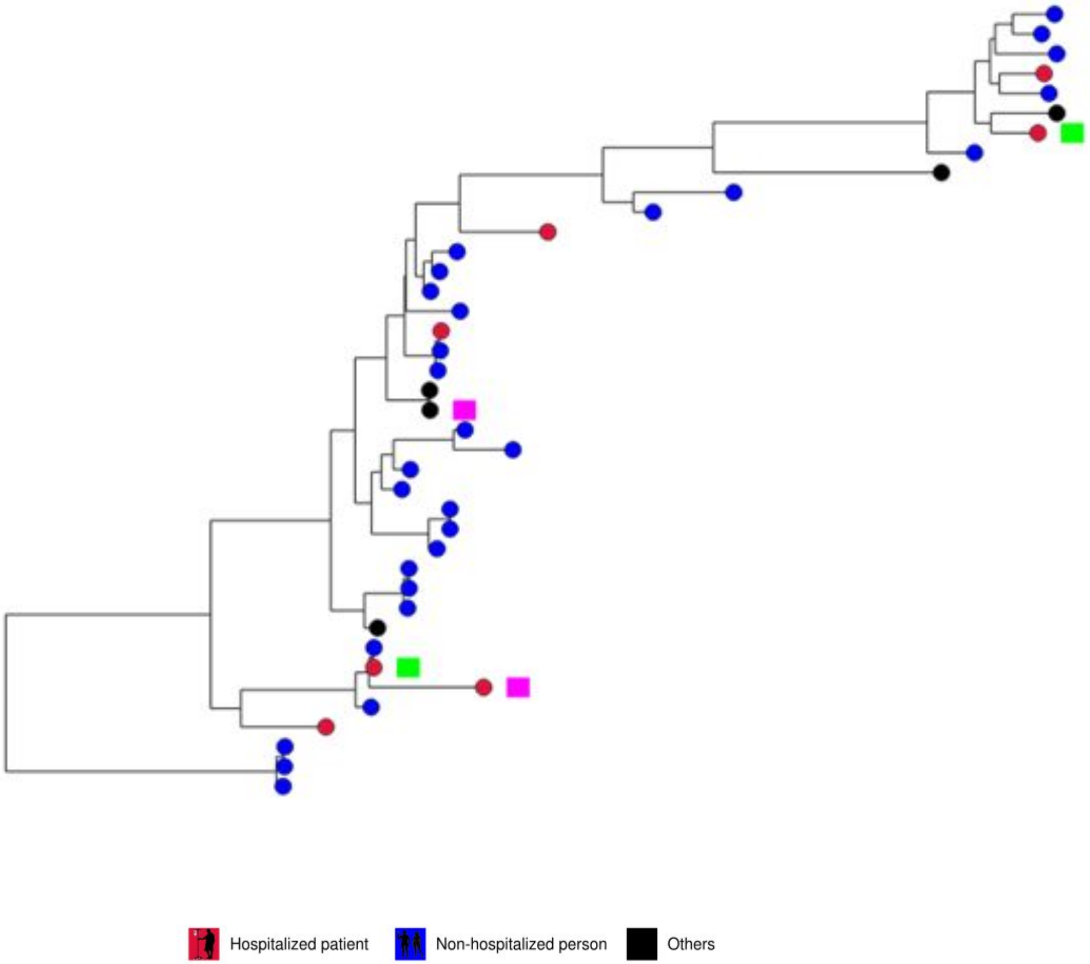


Supplemental Fig. S1. RAxML tree based on 859 core genes in 40 clade B isolates. Nodes represent the source with indication of presence of vancomycin resistance gene, vanA (pink) or vanB (green).

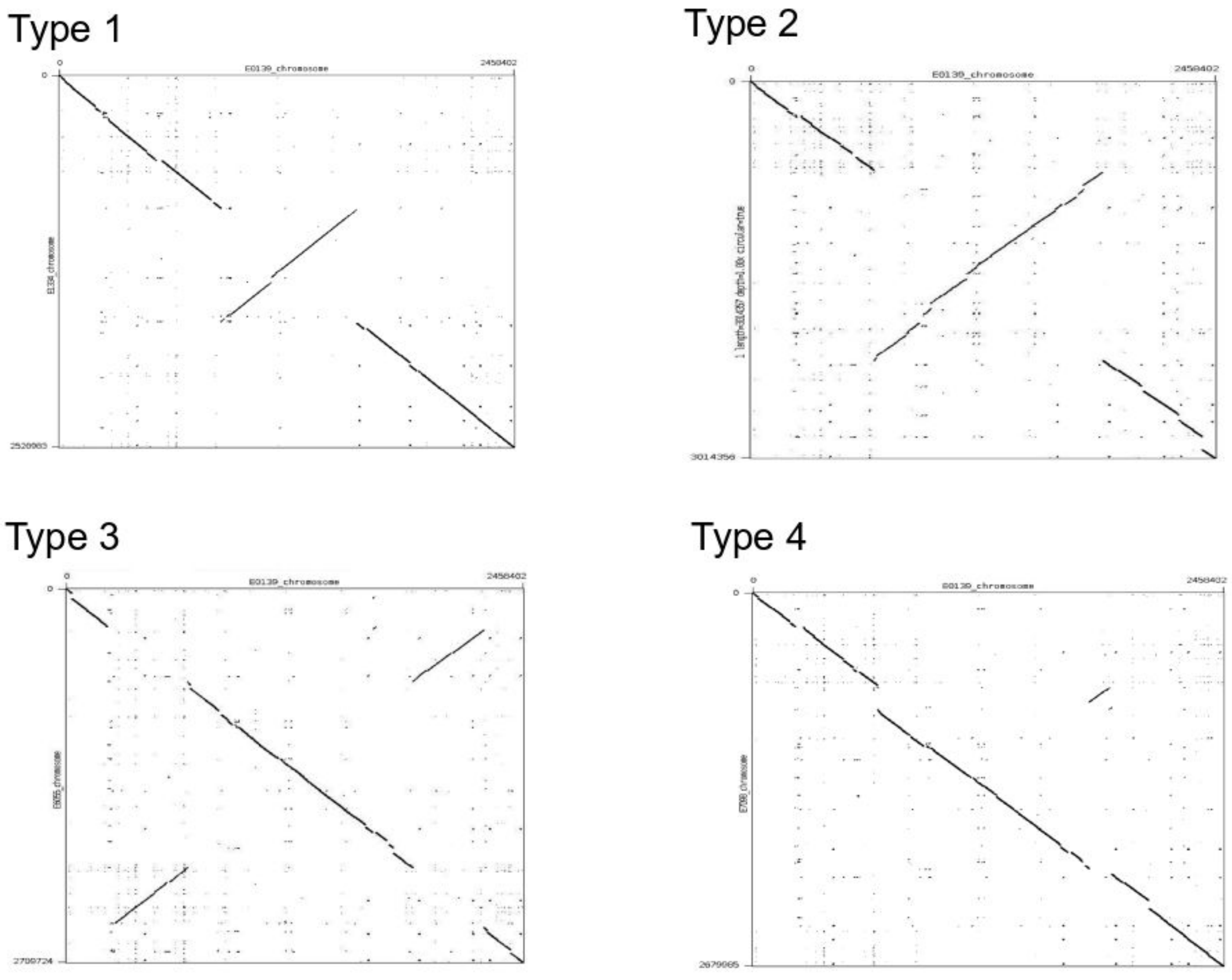


Supplemental Fig. S2. Dotplots of chromosomal rearrangements observed in our set of 48 complete chromosome sequences using strain E0139 (Clade A2) as reference. A, Chromosomal rearrangement Type 1 (E1334). B, Chromosomal rearrangement Type 2 (E8202). C, Chromosomal rearrangement type 3 (E6055). D, Chromosomal rearrangement type 4 (E7098).

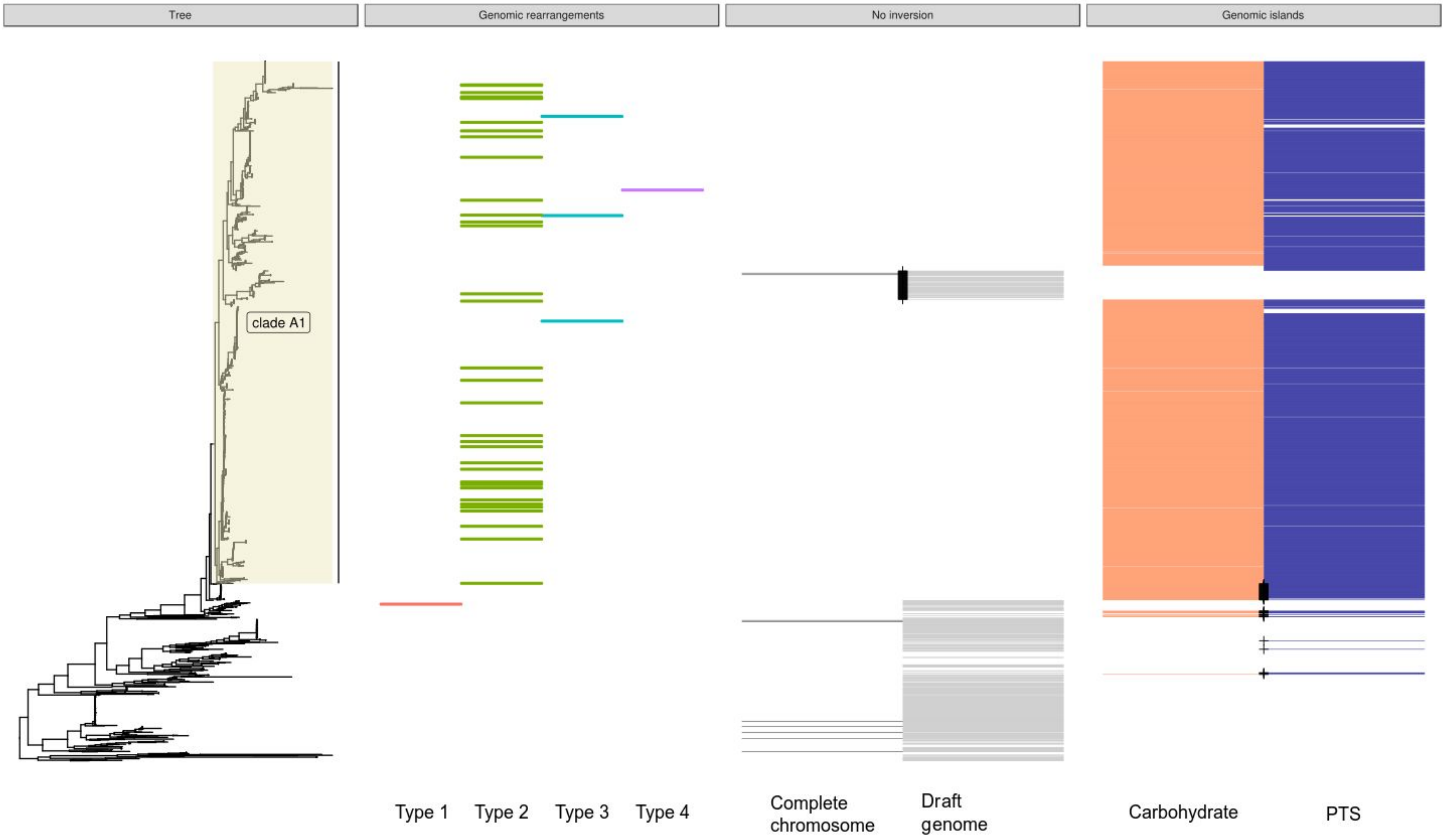


Supplemental Fig S3. Core genome tree based on 1,644 strains with the three metadata panels: i) distribution of genomic rearrangements types among 38 complete chromosomes, ii) indication of strains lacking a genomic rearrangement among 10 complete chromosomes and draft genomes and iii) indication of strains with insertion of a carbohydrate transport system encoding genomic island (orange)[33] or a phosphotransferase system encoding genomic island (purple)[34]. Black cross: indication of clade A1 dog isolates that lack the insertions/inversion or non-clade A1 dog isolates that do contain insertions/inversion.

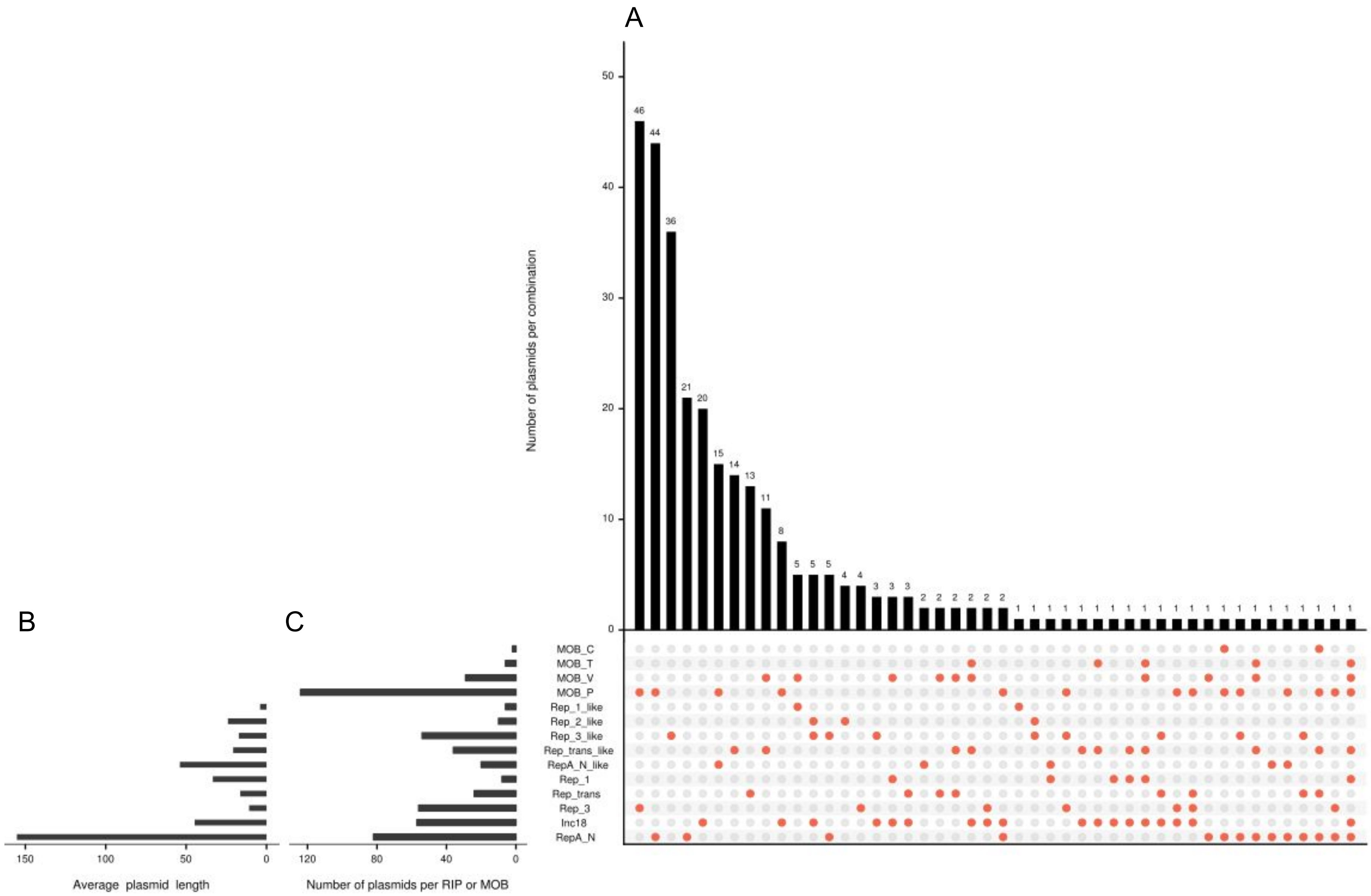


Supplemental Fig. S4. Overview of the complete plasmid sequences (n = 294) with an associated replication initiator gene (RIP). A) Histogram of the number of plasmids based on the combinations of RIP and MOB groups. B) Mean plasmid length (kbp) per RIP group. C) Number of plasmids per RIP or relaxase (MOB) group.

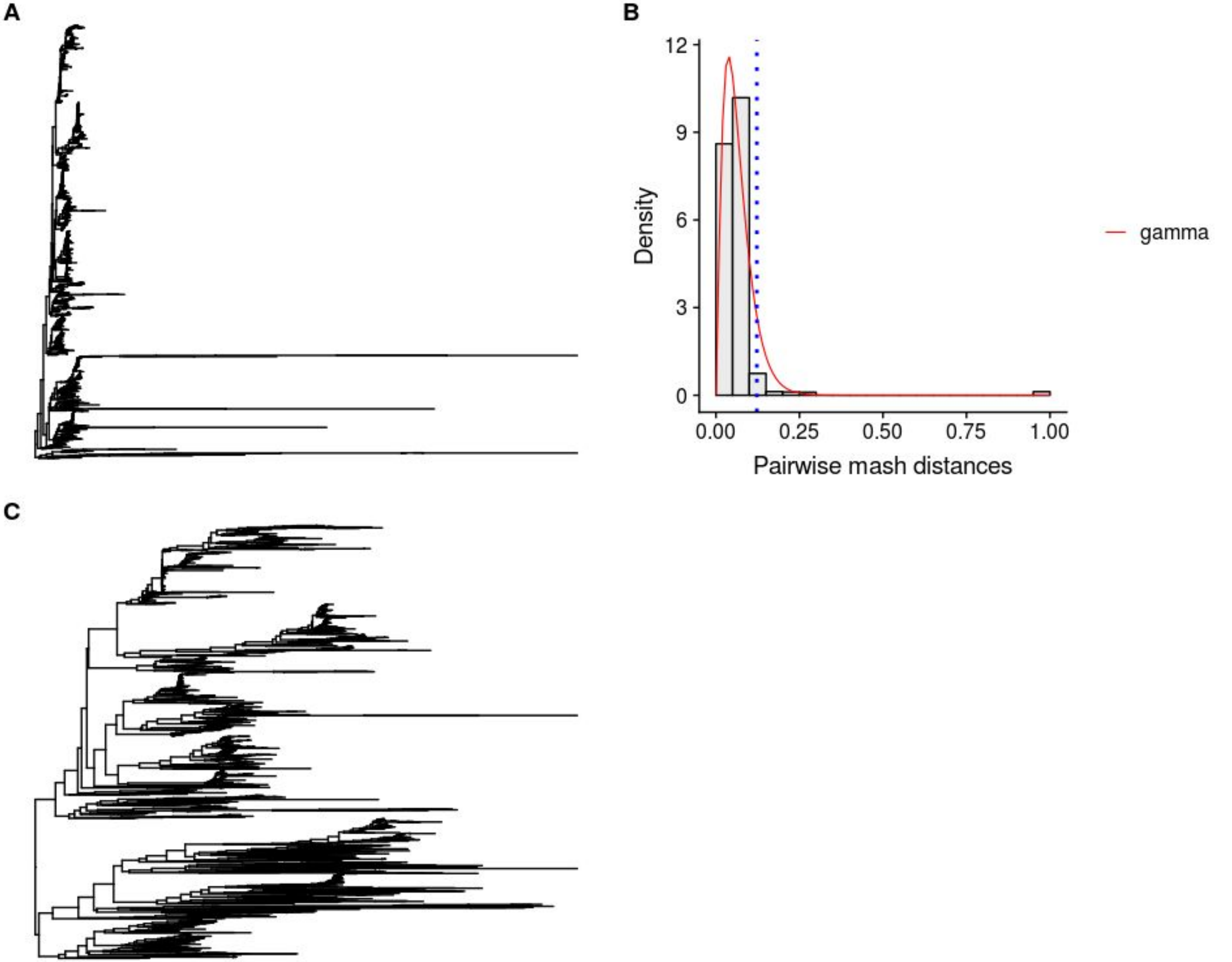


Supplemental Fig. S5. Maximizing resolution of bioNJ plasmid-based phylogeny. A) bioNJ tree of 1,639 isolates considering plasmid-predicted contigs by mlplasmids. B). Histogram against fitted density functions of pairwise Mash distances obtained by denscomp function (fitdistrplus R package). Vertical dash line indicates the Mash distance (0.1224967) used to filter out isolates (n = 37) with a higher average pairwise Mash distance. C) bioNJ plasmid-based phylogeny of 1,607 isolates after exclusion of 32 isolates.

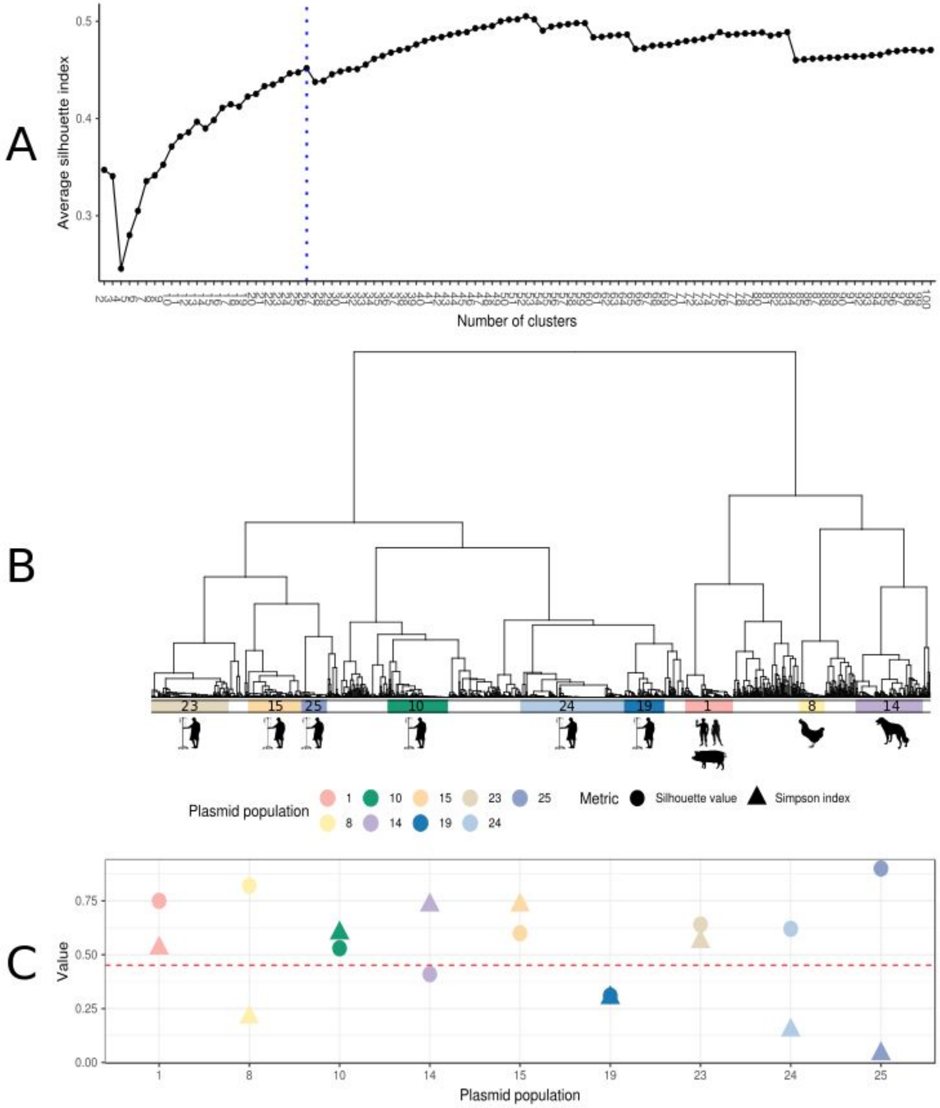


Supplemental Fig. S6. Definition of plasmid populations. A) Plasmid dissimilarity matrix of pairwise Mash distances was clustered using hierarchical clustering (ward.D2). Average silhouette index was computed based on the number of clusters (2-100). Vertical dash line corresponds to the clustering solution (k = 26) selected to cut the resulting dendrogram into 26 clusters. B) From these 26 clusters, we only defined plasmid populations (n = 9) if a particular cluster had a size larger than 50 isolates and an average silhouette index higher than 0.3. Each plasmid population was overrepresented at least with one isolation source. C) Simpson index (based on diversity of SC groups) and average silhouette index of each plasmid population. Horizontal dashed line indicates the average silhouette index of the selected clustering solution (k = 26, avg. silhouette index = 0.42).

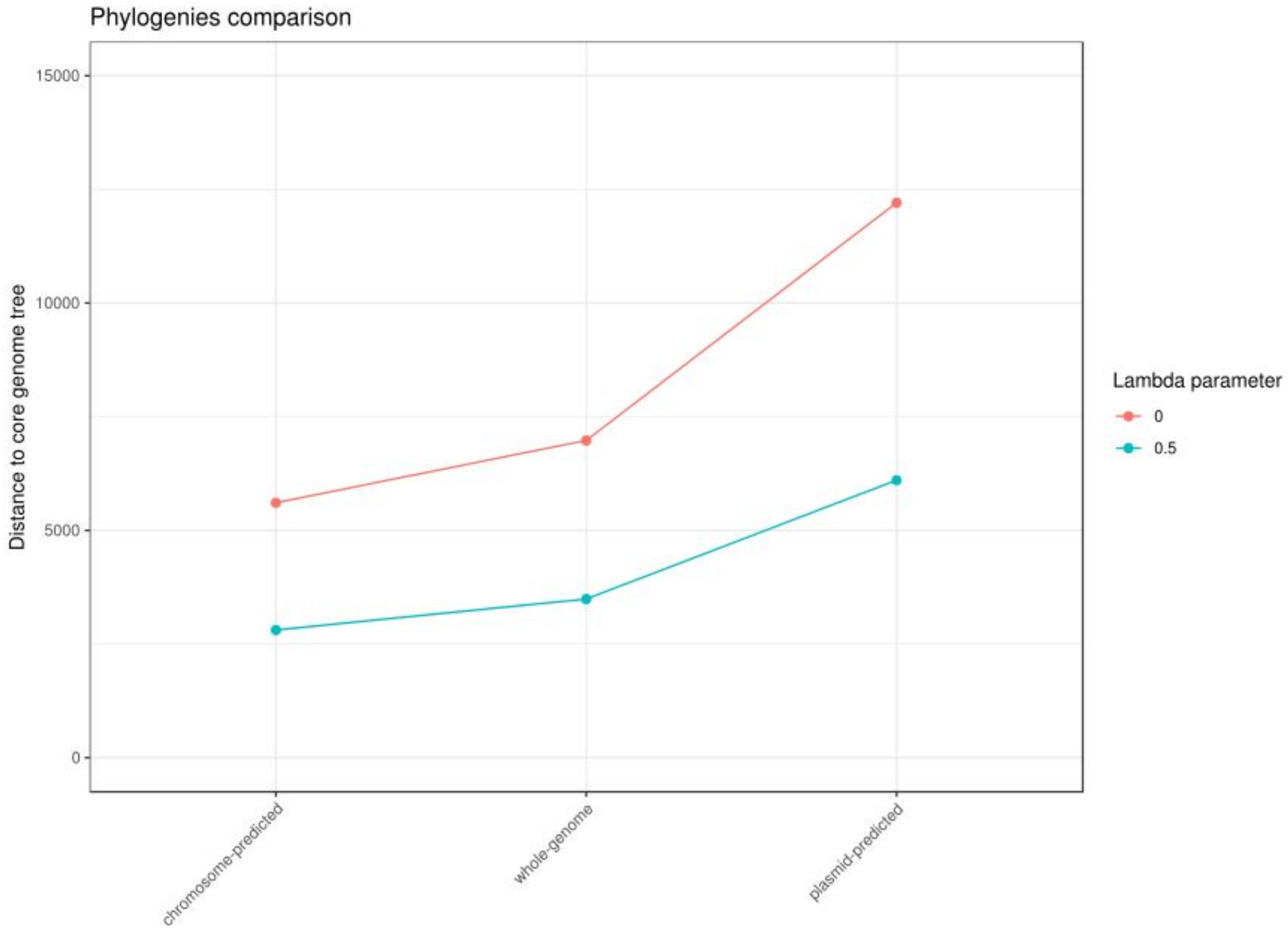


Supplemental Fig. S7. Phylogenies comparison (chromosome-predicted, whole-genome and plasmid-predicted) based on Kendall-Colijn (KC) metric considering differences only between topology of the trees ($\lambda$ = 0) and a balanced measure between branch and topology of the trees ($\lambda$ = 0.5) using as ground truth the core genome alignment.

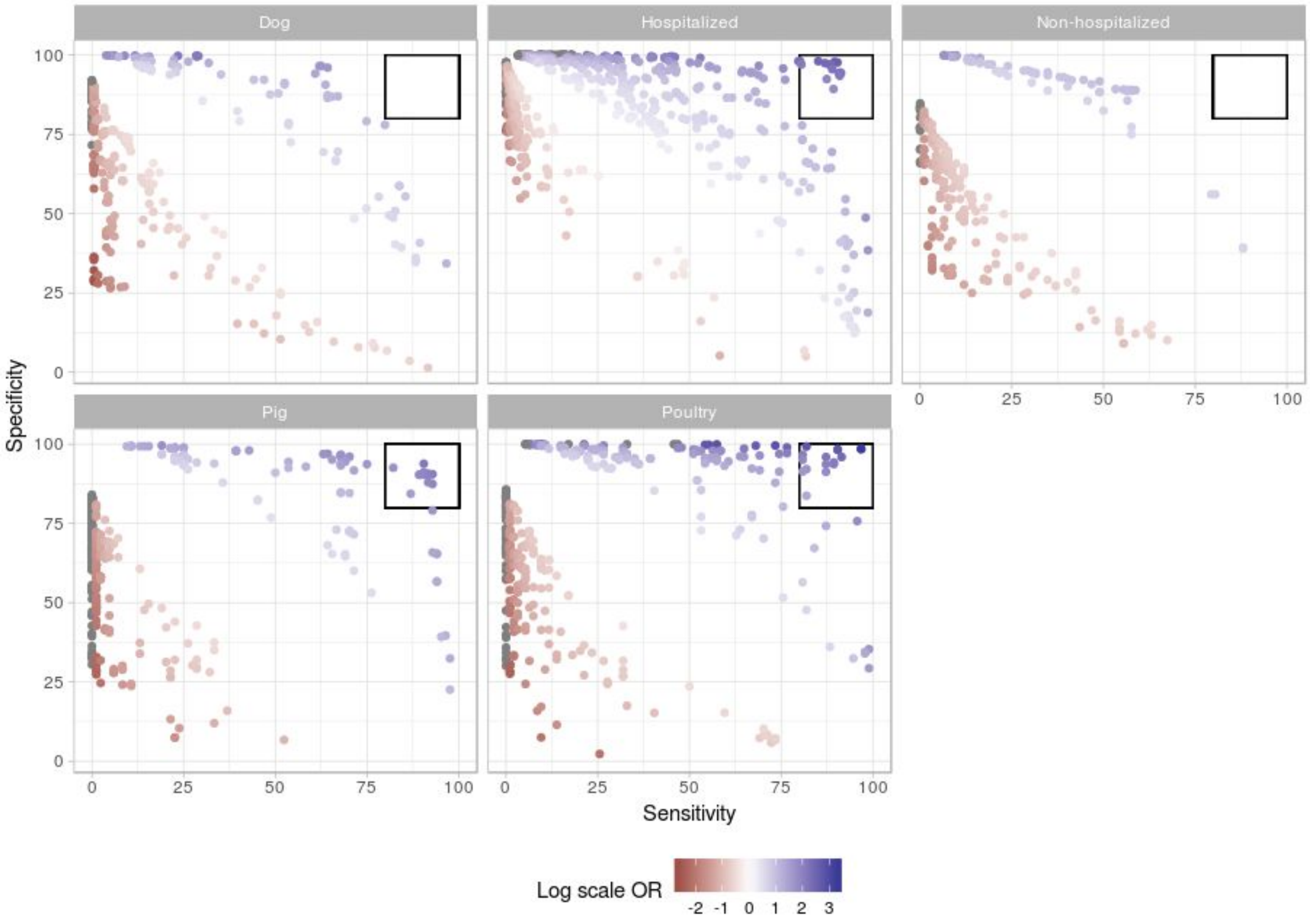


Supplemental Fig. S8. Distribution of orthologous genes (OG) groups based on isolation source. For each isolation source, Odds-ratio (OR) was transformed into a log scale (-2 to 3) to colour OG groups underrepresented (red) and overrepresented (blue). We further characterized OG groups (black square) with a specificity and sensitivity higher than 80%.

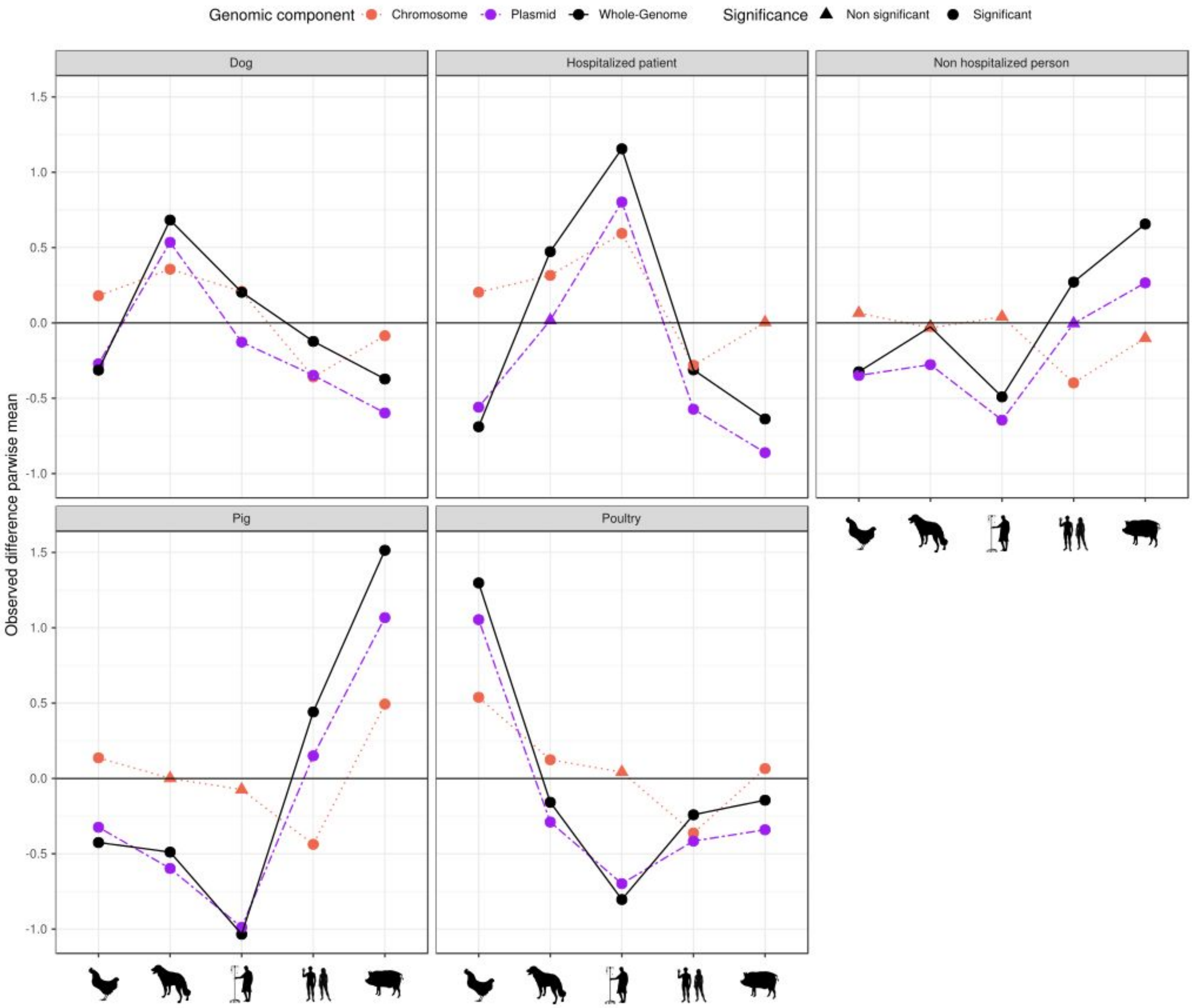


Supplemental Fig. S9. Differences in the observed means of average pairwise distances when comparing within-host and between-host groups against our defined random group of isolates. Each line corresponds to a different genomic component (whole genome - black solid, chromosome - dotted red, plasmid - dashed purple) and test significance is indicated based on shape (triangle - non significant, circle - significant).

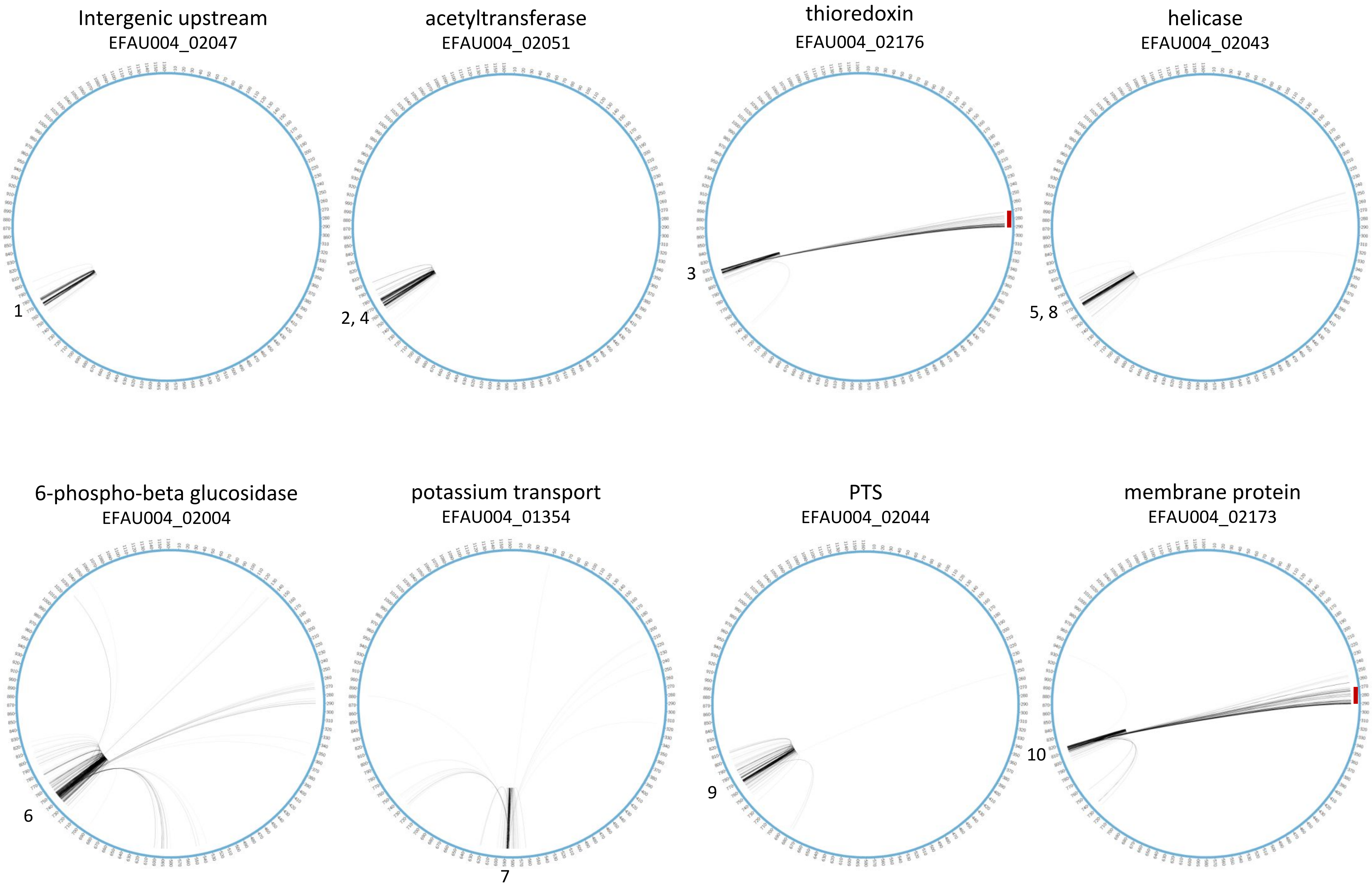


Supplemental Fig. S10. Circos plots of the core genome representing linked positions for ten loci with the highest number of linked SNPs. At the top of each plot, the converted position of loci in the AUS0004 genome.
